# High-content phenotypic analysis of *a C. elegans* recombinant inbred population identifies genetic and molecular regulators of lifespan

**DOI:** 10.1101/2024.01.15.575638

**Authors:** Arwen W. Gao, Gaby El Alam, Yunyun Zhu, Weisha Li, Elena Katsyuba, Jonathan Sulc, Terytty Y. Li, Xiaoxu Li, Katherine A. Overmyer, Amelia Lalou, Laurent Mouchiroud, Maroun Bou Sleiman, Matteo Cornaglia, Jean-David Morel, Riekelt H. Houtkooper, Joshua J. Coon, Johan Auwerx

## Abstract

Lifespan is influenced by complex interactions between genetic and environmental factors. Studying those factors in model organisms of a single genetic background limits their translational value for humans. Here, we mapped lifespan determinants in 85 genetically diverse *C. elegans* recombinant intercross advanced inbred lines (RIAILs). We assessed molecular profiles – transcriptome, proteome, and lipidome – and life-history traits, including lifespan, development, growth dynamics, and reproduction. RIAILs exhibited large variations in lifespan, which positively correlated with developmental time. Among the top candidates obtained from multi-omics data integration and QTL mapping, we validated known and novel longevity modulators, including *rict-1*, *gfm-1* and *mltn-1*. We translated their relevance to humans using UK Biobank data and showed that variants in *RICTOR* and *GFM1* are associated with an elevated risk of age-related heart disease, dementia, diabetes, kidney, and liver diseases. We organized our dataset as a resource (https://lisp-lms.shinyapps.io/RIAILs/) that allows interactive explorations for new longevity targets.

## Introduction

An intricate interplay of genetic, epigenetic, and environmental factors collectively determines the lifespan of an organism (Govindaraju et al., 2015). Over the past few decades, extensive research has been carried out to decipher the underlying mechanisms governing longevity with gain- and loss-of-function (G/LOF) studies in different model organisms. However, a prevalent limitation in the evaluation of the effects of mutations and environmental perturbations is the predominant reliance on animal models with a single genetic background for analysis (Li and Auwerx, 2020). This restricts the translational value and generalizability of these studies (Nadeau and Auwerx, 2019; Williams and Auwerx, 2015). Although such a strategy should ideally be employed in vertebrate models, the scope of the experimental testing in multiple genetic backgrounds combined with the ethical hurdles associated with such massive animal experimentation make this approach unrealistic. To overcome these constraints, the roundworm *C. elegans* has emerged as an attractive model for aging research, offering one of the best compromises between the simplicity of cell models and the complexity of vertebrate models (Gao et al., 2018b). In this regard, worm genetic reference populations (GRPs), such as the recombinant inbred lines (RILs) (Gao et al., 2018a; Li et al., 2010; Rockman et al., 2010; Vinuela et al., 2010) and recombinant inbred advanced intercross lines (RIAILs) (Andersen et al., 2015; Rockman and Kruglyak, 2009), have been increasingly used in the past years. These panels consist of inbred strains derived from crosses between two genetically divergent parental strains (Andersen et al., 2015; Thompson et al., 2015). With this study design, the recombination between the parental strains allows for fine mapping of quantitative trait genes (QTGs) — genes that explain the variation in certain quantitative traits (Evans et al., 2021). Furthermore, the availability of genotype data, and the ability to reproduce identical individuals, allow for the in-depth interrogation of quantitative traits at the systems level in several environmental conditions and at multiple physiological levels.

Here, we used a worm GRP consisting of 85 genetically diverse RIAILs derived from crosses between two parental strains, i.e., QX1430 (with an N2 Bristol background) and CB4856 (Hawaii) (Andersen et al., 2015; Gao et al., 2022). To investigate the alleles contributing to subtle variations in longevity-related phenotypes across this worm GRP, we measured their transcriptome, proteome, lipidome, and lifespan (Figure 1). In addition, we employed a high throughput fully automated microfluidic-based robotic phenotyping platform (see Nagi Bioscience SA https://nagibio.ch/), to collect other life-history phenotypes including body size, developmental dynamics, activity, as well as parameters related to worm reproduction and fertility. Integration of these omics and phenotypic data allowed the identification of a genetic locus associated with lifespan variations in these RIAILs. Within these loci, we identified *gfm-1*, *rict-1* and *mltn-1* as candidate longevity regulators and further validated *gfm-1* and *mltn-1* as bona-fide longevity regulators through loss-of-function studies. To assess the clinical significance of these candidate longevity genes in humans, we explored the UK Biobank data to show that variants in the human *GFM1* and *RICTOR* genes correlate with a variety of age-related disorders, such as heart disease, dementia, diabetes, kidney failure, liver disease and death. While our study focused on longevity regulation, we generated an extensive map of the molecular and phenotypic landscape in the RIAILs population. This resource will be valuable for subsequent *in silico* hypothesis generation and we have made it publicly available through an interactive open-access web resource (https://lisp-lms.shinyapps.io/RIAILs/).

**Figure 1.**
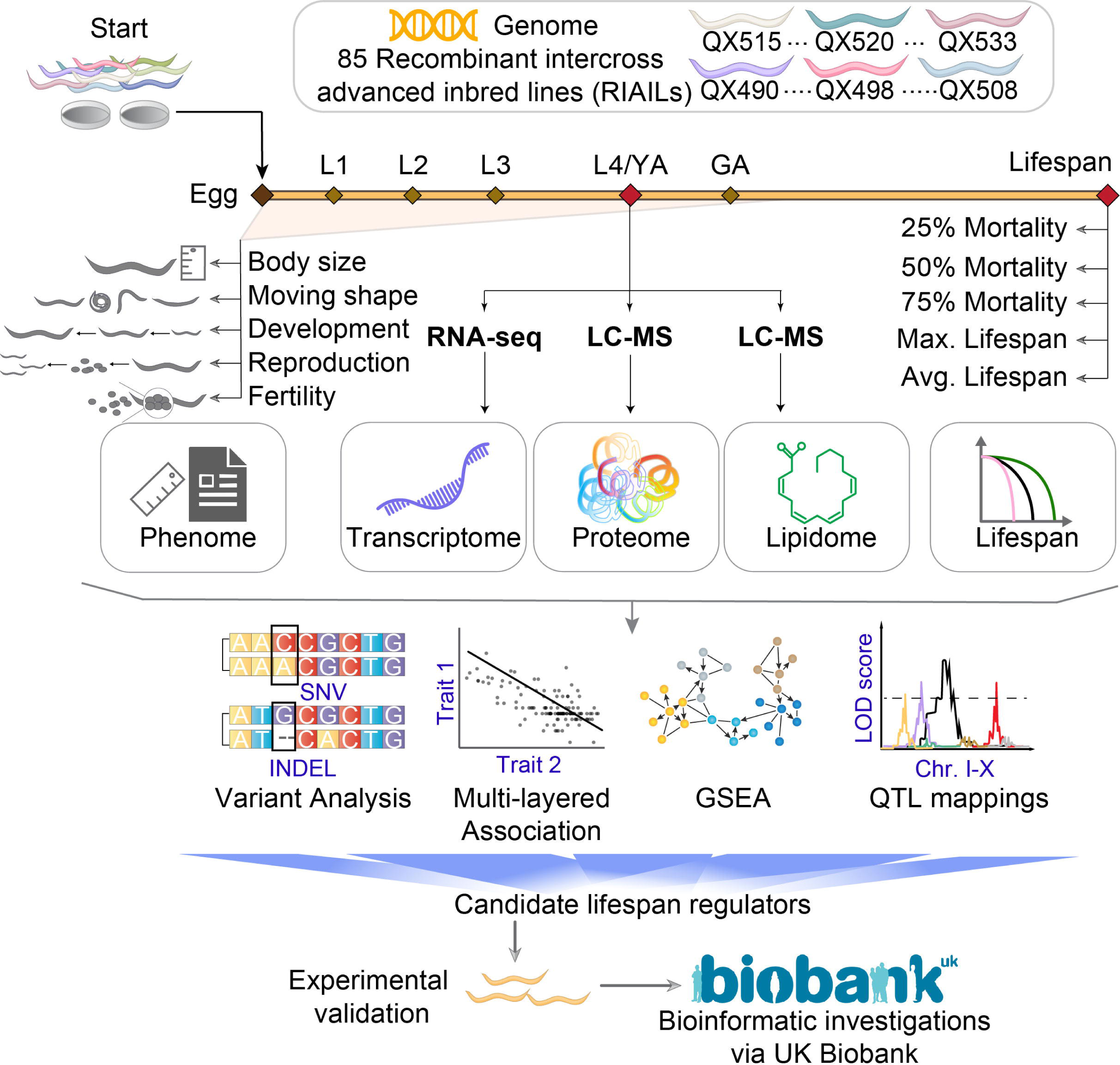
Overview of the study design. 85 recombinant intercross advanced inbred lines (RIAILs) derived from the crossing of QX1430 (N2 Bristol background, with deletions of confounder genes) and CB4856 strain (Hawaii) were used. Lifespan, early life-history traits, transcriptome, proteome, and lipidome were collected for each strain. We applied a systems genetics approach to study relations between different phenotypes and molecular traits to identify candidate lifespan genes. After prioritization of the candidate genes, we validated them through wet lab experiments and human population genetics (e.g. UK Biobank). We collected data from all RIAILs using three pipelines. In the first, worms were cultured and scored for their lifespans; in the second, they were cultured in a microfluidic device for ∼100 h to collect early life-history traits, including body size, moving shapes, developmental parameters, reproduction, and fertility; and in the last, they were cultured to reach L4/young adulthood, and collected for multi-omics measurements (transcriptomics, proteomics, and lipidomics). Examples of variants: SNV: single-nucleotide variant. INDEL: insertions and deletions. GSEA: gene set enrichment analysis; Chr. I-X: chromosome I to X; LC-MS: liquid chromatography–mass spectrometry; L1-L4: larval stage 1 to 4; YA: young adulthood; GA: gravid adulthood. Max. Lifespan: maximum lifespan; Avg. Lifespan: average lifespan.

## Results

### 85 RIAILs exhibit extensive variations in the lifespans and life-history traits

To determine the extent to which genetic background can influence longevity, we first assessed the lifespans of 85 RIAILs by manually scoring them on plates as described previously (Gao et al., 2022). The range of average lifespan of RIAILs was from 13 to 21 days (Figure 2A). Although the majority of RIAIL average lifespans lay between those of the parentals, nine strains’ lifespans were shorter lived than CB4856 and four strains lived longer than N2, suggesting the presence of transgressive segregation (Figures 2A and 2B). Also, we observed a similar pattern for early (25% dead and 75% alive), mid (50% mortality), and late (75% dead and 25% alive) time to mortality (Figure 2C), that is, the age in days at which 25%, 50%, or 75% of the worms died, respectively. Studies in different organisms have shown that diverse trade-offs dominate life-history traits (Blueweiss et al., 1978; Maklakov and Chapman, 2019; Thomas Flatt (ed.), 2011). As a consequence, various organisms display correlations among different life history traits, such as lifespan and fecundity (Luckinbill et al., 1984), development and lifespan (Marchionni et al., 2020), body size and longevity (Blanco and Sherman, 2005; Bou Sleiman et al., 2022; de Magalhaes et al., 2007). We therefore monitored a number of life-history traits across the early life stage of the RIAILs (approximately 100 h after egg hatching), including maximum body size, developmental time, sexual maturity (emergence of the 1^st^ egg), fertility (rate of egg accumulation), embryonic viability (emergence of the 1^st^ larvae, following the emergence of the 1^st^ egg), and the rate of progeny accumulation, using an innovative whole-organism high-content screening technology (see https://nagibio.ch/) (Atakan et al., 2018) (Figures 2D-2E and S1). RIAIL strains displayed large variations in early life-history phenotypes, including developmental dynamics, reproduction (Figure 2D), and activity (Figure 2E). The N2 strain had a protracted growth period, characterized by delayed attainment of maximal body size and greater overall body size when compared to the CB4856 strain (Figures 2D and S1A). Both strains had comparable timing of reproductive maturation and showed no significant disparities in various fertility measures (Figures 2D and S1C-S1D).

**Figure 2.**
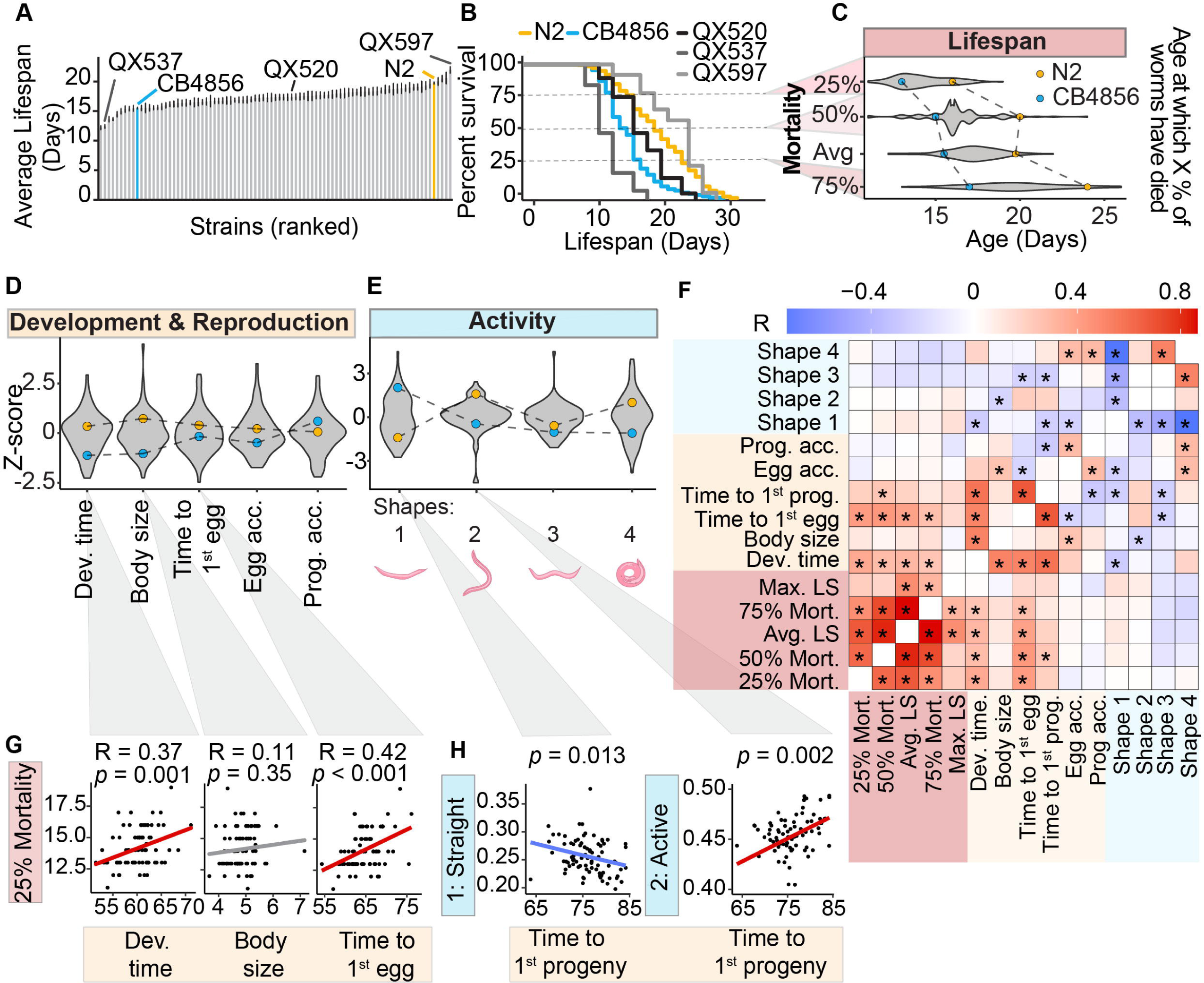
RIAILs exhibit extensive variation in lifespan and life-history traits. (**A**) Bar plot showing the average lifespan of 85 RIAIL strains (60 worms/strain) and two wild-type parental strains (600 worms/strain). Grey bars: RIAILs; Orange bar: N2 (Bristol); Blue bar: CB4856 (Hawaii, HW). Examples of strains with different average lifespans are labeled. (**B**) Examples of differences in the average lifespan of RIAILs (QX537, QX520, and QX597; grey) and parental N2 (orange) and CB4856 (blue) strains. (**C**) Violin plots of the RIAIL lifespan traits. Dots represent the average value of the trait for the two parental strains. (**D**) Violin plots of early life-history phenotypic traits. Dev. time: developmental time; Egg acc.: egg accumulation; Prog. acc.: progeny accumulation (**E**) Violin plots of the activity life-history phenotypic traits. Shape 1: straight; Shape 2: active; Shape 3: swimming; Shape 4: supercoil. (**F**) Pearson correlation between lifespan and physiological traits. Stars represent non-adjusted p-values (*: adjusted BH p-value < 0.05). LS: lifespan. Mort: mortality. (**G**) Correlation of 25% mortality with developmental time, body size, and the time to the 1^st^ egg, respectively. R: Pearson correlation coefficients *p*: p-values. (**H**) Scatter plots of time spent in 2 moving shapes and time to the 1^st^ progeny of each RIAIL strain. The p-value indicates the coefficient in the linear model. **Related to Figures S1 and S2.**

### Longer lifespan is associated with slow development and late egg emergence

Early life-history traits can potentially provide insights into the developmental trajectory and long-term outcomes of organisms (Bou Sleiman et al., 2022; Miller, 2002). To obtain an overview of associations between the phenotypic traits and lifespan traits, we pooled phenotypes and performed pairwise correlation analysis between all traits (Figure 2F). We found that, worm developmental time (r^2^ = 0.37, p < 0.001), reflecting the worm growth dynamics, and egg emergence (time to the moment when a worm lays its first egg, r^2^ = 0.42, p < 0.001), reflecting sexual maturity, were the most associated with 25% mortality (Figures 2F-2G). Both body size and progeny emergence (time until the first progeny of a worm is detected) were strongly correlated with developmental time (r^2^ = 0.52, p < 0.001 and r^2^ = 0.58, p < 0.001 respectively). In addition to the phenotypic readouts on worm development and reproduction, we also evaluated the most common shapes of worms in each population (Figures 2E and S2A). Four main categories of shapes were defined as follows: two regular wild-type shapes in liquid (shape 2 - active; shape 3 - swimming) and two extreme shapes (shape 1 - straight; shape 4 - supercoiled) (Figure 2E). Since worms adopt different shapes over time, the shape metric reflects the percentage of time worms spent in each shape category. We observed a negative association between shape 1 (straight) and the duration before the first progeny appeared. Conversely, there is a positive correlation between shape 2 (active movement) and the time it took for the first progeny to emerge (Figures 2F and 2H). However, none of the shapes directly correlated with the lifespan traits (Figures 2F and S2B). In combination, these findings corroborate that early life-history traits, such as delayed development, are an evolutionary cost of longevity.

### Early life transcriptome unveils potential pathways influencing lifespan traits

To explore connections between the transcriptome at the early life (L4) stage and longevity, we performed an association analysis between the expression levels of ∼20,000 transcripts and three measures of time to mortality: 25%, 50%, and 75% (Figures 3A and S3). Notably, a substantial proportion of these transcripts exhibited significant associations (unadjusted p-values) with time to 25% mortality, while associations with those of 50% and 75% time of mortality and average lifespan were less pronounced (Figure 3B). Specifically, 1,951 transcripts correlated significantly with 25% mortality (1,028 positives and 923 negatives), while 894 transcripts were significantly linked to 50% mortality (636 positives and 258 negatives), and 189 transcripts showed significant associations with 75% mortality (109 positives and 80 negatives) in the RIAIL population (Figure 3C). The predominant correlation of transcripts with early mortality might stem from the fact that the transcript profiles were extracted from L4/young adult worms. Additionally, 504 transcripts were nominally significantly associated with the average lifespan in RIAILs (Table S1). None of the associations between transcripts and lifespan traits remained significant after multiple testing corrections; therefore, we explored the possibility of significant disparities at the pathway level using gene set enrichment analysis (GSEA) to find out if there is any collective impact of the transcripts (Figure 3D). We found 938 pathways significantly enriched for 25% mortality (81 positive and 857 negative), 701 (105 positive and 596 negative) for those associated with 50% mortality, and 58 (32 positive and 26 negative) for 75% mortality respectively (Figure S3 and Table S2). Given the early life transcriptome was more strongly associated with 25% mortality than other lifespan traits (Figure 3B), we investigated the enriched pathways associated with this metric. The majority of 25% mortality-enriched biological processes were negatively associated with lifespan and were primarily involved in chromosome organization, cytoskeleton organization, cellular lipid metabolism, cell division, DNA repair, and protein metabolic processes (Figure 3D). Among the top 30 enriched pathways, two pathways were positively associated with 25% mortality, namely neuropeptide signaling and G protein-coupled receptor signaling. Although it was among the top 30, the geneset “determination of adult lifespan” was among those significantly inversely associated with early mortality (q-value < 0.01) (Figure 3E and Table S2). Lower expression of genes within this geneset was associated with a longer time to reach 25% mortality (Figure 3F). Taken together, the early life transcriptome showed significant associations with lifespan at the pathway level, particularly with 25% mortality. Our data indicate that various biological processes and pathways can influence early mortality and potentially affect longevity.

**Figure 3.**
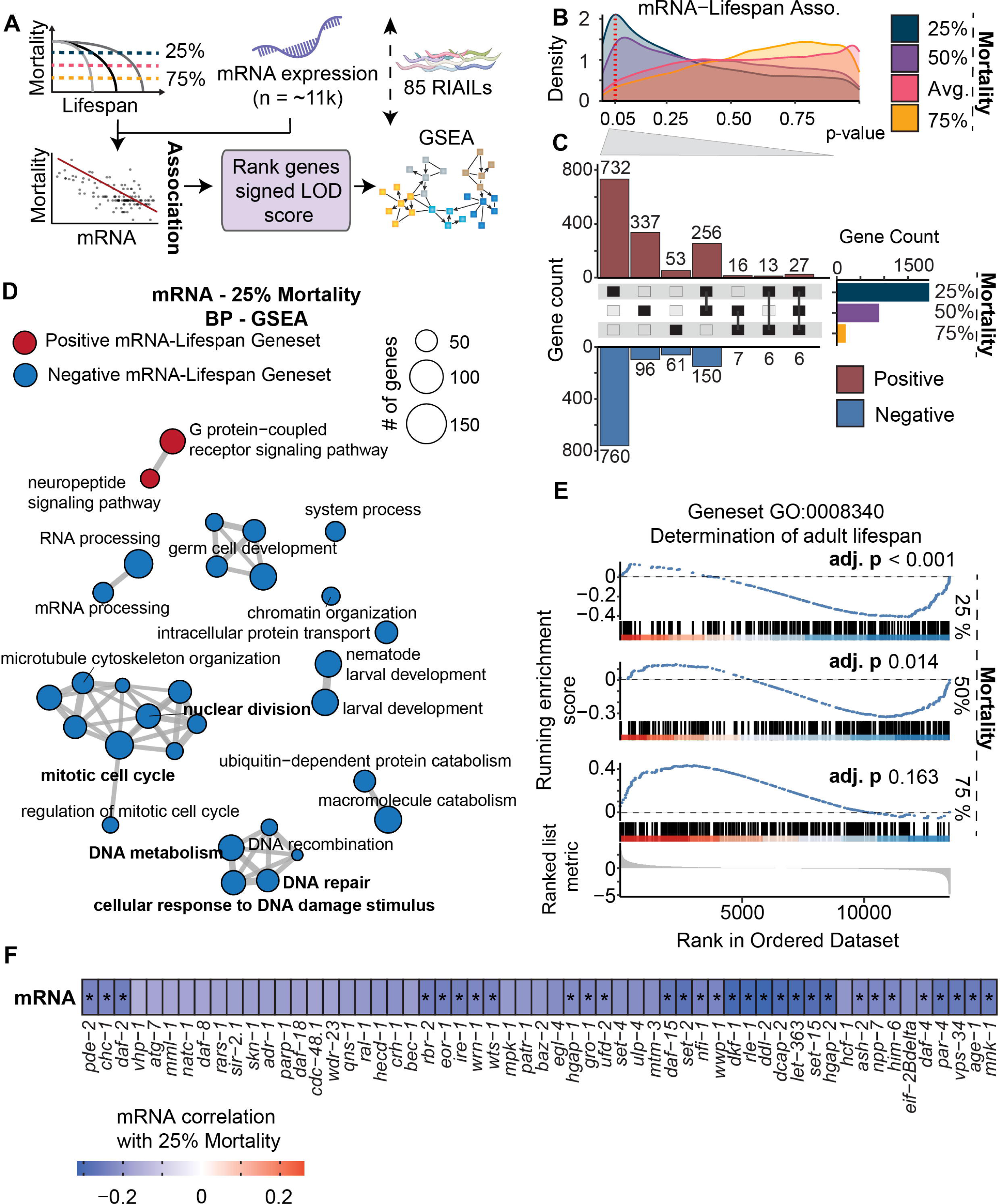
Quantitative assessment of transcriptome-lifespan associations. (**A**) Schematic pipeline of mRNA-lifespan association mapping and gene set enrichment analysis (GSEA). (**B**) Line histogram of non-adjusted p-values of mRNA-lifespan associations for the different lifespan traits. The distribution of mRNA-lifespan associations showed a higher density of significant associations (non-adjusted p-values) for the 25% mortality trait. The Y-axis represents the density of p-values. The red dashed vertical line represents a p-value of 0.05. Color represents the different lifespan traits. (**C**) The upset plot of positively-(red) and negatively-(blue) associated genes with 25%, 50%, and 75% mortality in the RIAILs with a non-adjusted p-value smaller than 0.05. (**D**) Graph representing the top 30 biological process gene sets enriched. GSEA of mRNA-25% mortality associations. Genes were ranked by the signed logarithm of the odds (LOD) score of mRNA-25% mortality association. Color represents positive (red) or negative (blue) normalized enrichment score (NES). All gene sets in the figure had a significant q-value. Genesets in bold: overlapping genesets in both mRNA and protein levels. (**E**) The running GSEA plot of “determination of adult lifespan” (GO:0008340). adj. p: adjusted p-value. (**F**) Heatmap of Pearson correlation of transcripts enriching for GO:0008340 with 25% mortality in the RIAILs. Star represents a non-adjusted p-value < 0.05. **Related to Figure S3 and Tables S1 and S2.**

### Quantitative assessment of correlations between protein pathways and different lifespan traits

As proteome analysis offers a more direct perspective on cellular function, complementing the information obtained through transcriptome analysis, we measured the protein profiles of RIAILs and detected >6,500 proteins following the removal of non-detectable peptides and rigorous quality control measures (Figure S4A and Table S3). In contrast to the mRNA-lifespan association analysis, the number of proteins showing significant association with 25% mortality (non-adjusted p-values) was comparable to those linked with 50%, 75% mortality, and the average lifespan (Figures S4B-S4C). When investigating the top pathways enriched for 25% mortality, we found seven pathways, including vesicle-mediated transport, Golgi vesicle transport, actomyosin structure organization, and supramolecular fiber organization, that positively correlated with the 25% mortality (Figure S4D and Table S3). The negatively associated pathways were mostly related to DNA metabolism and cell cycle regulation (Figure S4D and Table S3). In addition, we examined the pathways enriched at both the mRNA and protein levels (Figures 3D and S4D). Gene sets involved in DNA damage response, DNA repair, and cell cycles overlapped at both mRNA and protein levels and were negatively correlated with lifespan traits. Consistent with these results, cell cycle, and associated genome integrity pathways were reported to be negatively associated with cellular turnover, a measure of cell and tissue longevity (Seim et al., 2016), whereas enhanced DNA repair capacity has been suggested in long-lived species (Cortopassi and Wang, 1996; Ma et al., 2016).

### Cardiolipins (CLs), triglycerides (TGs), phosphatidylinositol (PIs) were positively and phosphatidylethanolamines (PEs) were negatively correlated with lifespan traits

Perturbations in circulating lipid levels due to genetic, lifestyle, and environmental factors can heighten the risk of developing age-related disorders, such as cardiovascular and metabolic diseases (Harshfield et al., 2021). To determine possible links between lipids and lifespan, we integrated lifespan traits with lipid profiles measured in the RIAIL cohort (Figure 4A). The first two dimensions of a principal component analysis (18.2% and 11.7% of variance explained, respectively) did not visually segregate strains by lifespan (Figure 4B). We then examined lipid-lifespan correlations (unadjusted p-values of less than 0.05) and highlighted distinct correlation profiles between lipids and different lifespan metrics (Figures 4C and 4D). Cardiolipins (CLs), comprised mainly of polyunsaturated acyl chains, were among the lipids that displayed a positive correlation with average lifespan and 75% mortality (Figure 4C and Table S4). The levels of CLs consistently decline in aged worms and rats (Gao et al., 2017; Smidak et al., 2017), supporting the concept that higher levels of CLs may be advantageous for health and longevity. Phosphatidylinositols (PIs) were among the primary lipid classes positively associated with average lifespan and 75% mortality, while many triglycerides (TGs) correlated positively with 25% mortality (Figure 4C). In contrast, numerous phosphatidylethanolamines (PEs) and PE-derivatives (e.g. plasmanyl-PE and plasmenyl-PE) exhibited a negative correlation with all the lifespan traits (Figure 4D and Table S4). Over-representation analysis of the different lipid classes confirmed that TGs, CLs, and PIs were positively associated with lifespan traits, while PEs were negatively associated (Figure S5).

**Figure 4.**
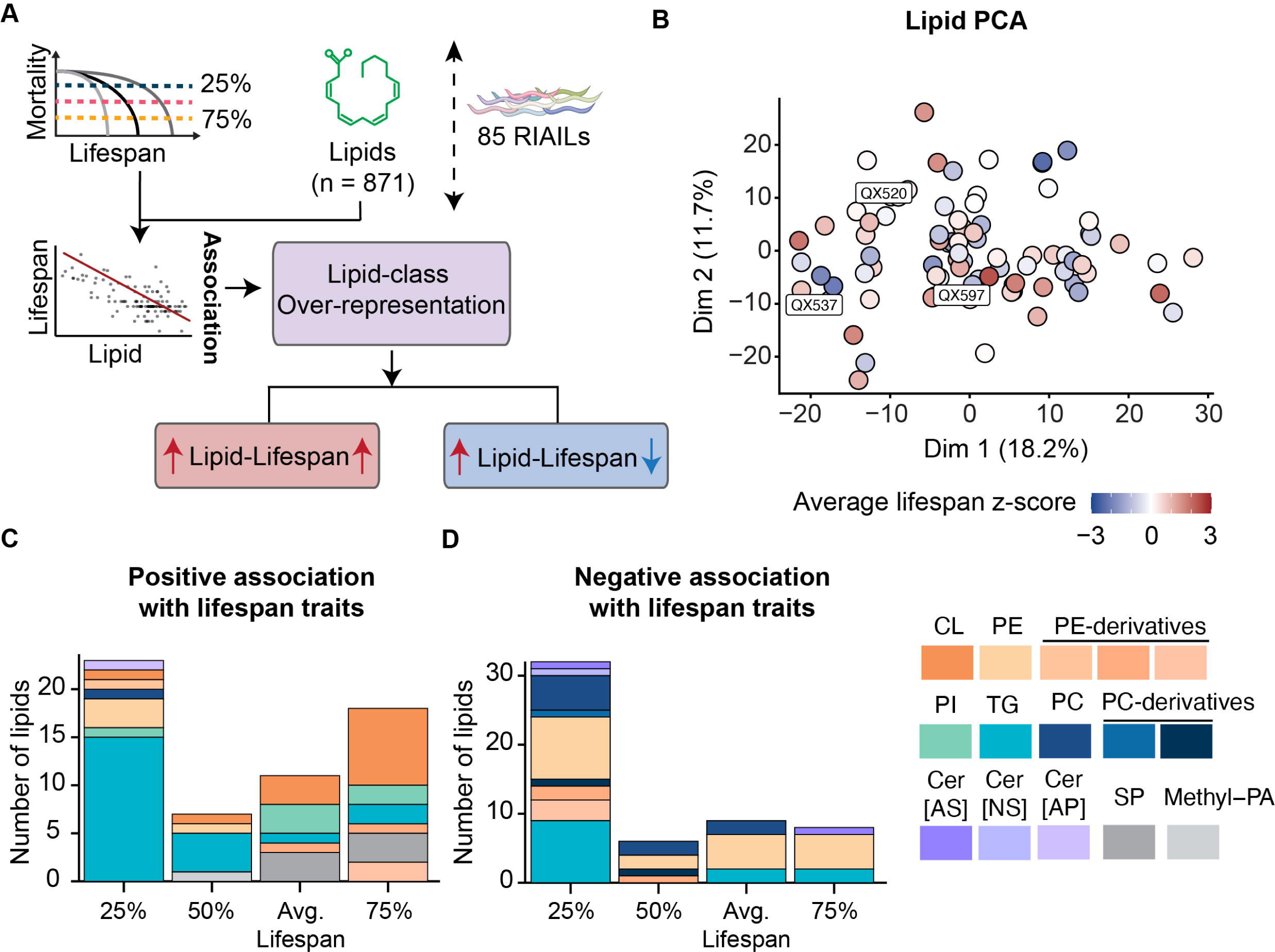
Quantitative assessment of lipidome-lifespan associations. (**A**) Diagram of lipid-lifespan association mapping and lipid-class over-representation. (**B**) Principal component analysis (PCA) representation of RIAIL strains based on all LC-MS measured lipids at L4/young adulthood. Color represents z-score of the average lifespan of each RIAIL strain. (**C-D**) Bar plots of the number of lipids with significant association (non-adjusted p-value < 0.05) and positive (C) or negative (D) association coefficient. CL: cardiolipin. PE: phosphatidylethanolamine. PE-derivatives: including Lyso-PE, plasmanyl-PE, plasmenyl-PE. PI: phosphatidylinositol. TG: triglycerides. PC: phosphatidylcholines. PC-derivatives: PC[OH] and plasmanyl-PC. Cer[AS]: ceramideAS. Cer[NS]: ceramideNS. Cer[AP]: ceramideAP. SP: Sphingolipid. Methyl-PA: methylphosphatidic acid. **Related to Figure S5 and Table S4.**

### Lifespan variations are not dependent on overall genomic composition nor mitochondrial haplotype

As the parental strain CB4856 worms have a shorter lifespan compared to the other parental strain N2 (Figures 2A-2B) (Gao et al., 2022), we asked whether the allelic proportion of each parental genome in the RIAILs could partly explain the variations observed in the lifespan traits. While we observed a considerable variation in parental allele distributions across the RIAIL genomes (Figure S6A), the phylogenic tree based on genetic distance between the strains did not cluster strains according to their lifespan (Figure S6B). For instance, strains QX580 and QX594 are highly genetically related (Figure S6B, next to the N2 strain), yet QX580 has a relatively longer lifespan in comparison to QX594, which displays a relatively shorter lifespan. The lifespan of CB4856 was previously found to be significantly influenced by variants in the mitochondrial DNA (mtDNA) (Dingley et al., 2014). We therefore separated the RIAILs by their mitochondrial genotypes but found no significant associations between the mitotype and the average lifespan (Figure S6C). A previous study using a different, small set of RIAILs found a positive correlation between the CB4856 mitotype and lifespan, as well as a negative correlation between N2 mitotype and lifespan (Zhu et al., 2015). However, we found no such correlation in either the CB4856 or N2 mitotype background in these RIAILs (Figure S6D). Taken together, these data underscore the necessity for a more refined approach to pinpoint specific loci that determine lifespan.

### Identification of a lifespan QTL on Chromosome II

Next, we sought to leverage the genetic diversity of the RIAIL population to map their associations with lifespan traits and potentially uncover novel genetic regulators of lifespan. Through variant calling using RNA-seq data (Figure S7), we generated a genetic map for the RIAIL population (Figure S8). We then performed quantitative trait loci (QTL) mapping of lifespan traits and detected a significant QTL on Chr. II for average lifespan (Figure 5A). We detected a suggestive QTL in the same locus for 25%,50%, and 75% mortality. We observed a decrease in the LOD score for this locus, from 4.13 for average lifespan to 3.83 for 50% mortality, and finally to 3.28 for 75% mortality (Table S5). Upon examining the average lifespan of the RIAILs for the two genotypes at this locus, we found that strains with the CB4856 genotype have longer lifespans compared to strains carrying the N2 genotype (Figure 5B). In other words, the allele associated with a longer lifespan comes from the shorter-lived CB4856 strain, suggesting that complex gene-gene interactions overcome any single-locus effect on lifespan in the RIAILs.

**Figure 5.**
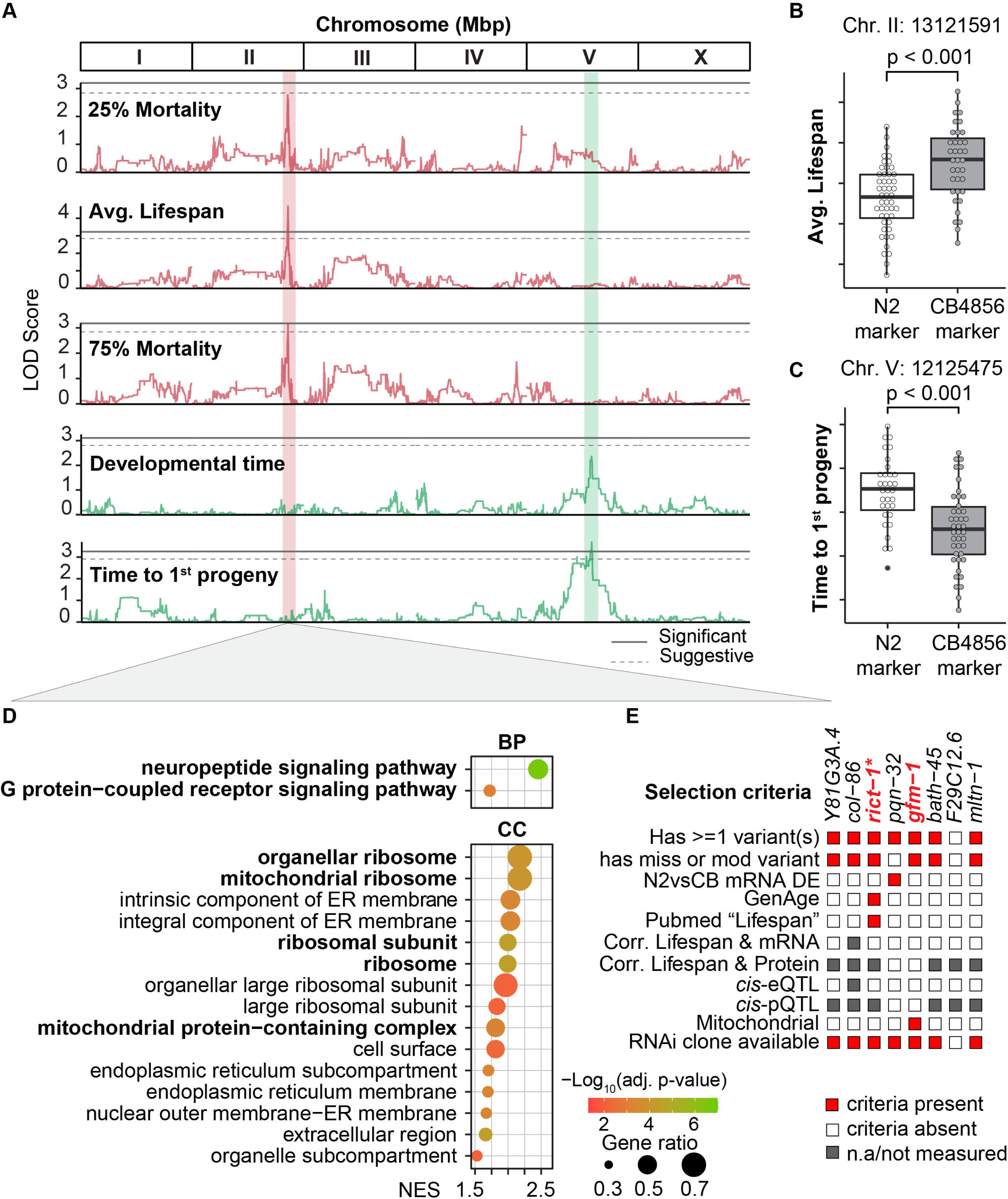
Identification of a lifespan-modulating locus on Chromosome II and prioritization of candidate genes. (**A**) QTL mapping of lifespan and life-history traits identifies two significant QTL on Chr. II, and V for average lifespan, and time to 1^st^ progeny, respectively. The vertical axis shows the logarithm of the odds (LOD score). The horizontal axis represents the genomic position in mega-basepair (Mbp). Dashed and solid grey lines represent suggestive (p < 0.1) and significant (p < 0.05) thresholds, respectively. (**B**) Boxplot of the average lifespan of RIAILs with CB4856 or N2 genotype at position 13,121,591 on Chr. II. (**C**) Boxplot of time to 1^st^ progeny of RIAIL strains with CB4856 or N2 genotype at position 12,125,475 on Chr. V. The p-value represents the comparison of the two groups calculated using a two-tailed Student’s t-test. (**D**) Gene set enrichment analysis based on differential transcriptome analysis at position 13,121,591 on Chr. II. Only the top 15 significant and positively enriched genesets for biological process terms (**upper**) and cellular component terms (**lower**) are shown. Color represents −log_10_(adjusted p-value). BP: Biological process. CC: Cellular component. NES: normalized enrichment score. Interesting gene sets were highlighted in bold. (**E**) Candidate genes are prioritized under the confidence region of the loci on Chr. II. Genes under the lifespan QTL peak were annotated if: a gene has one or more variants in CB4856; a gene has one or more missense or modifier variants in CB4856; the gene is differentially expressed between N2 and CB4856 (absolute log_2_ (fold change) > 1 and adjusted p-value < 0.05); the gene has been annotated in GenAge (https://genomics.senescence.info/genes/index.html) for involvement in the aging; the gene has been found in a paper with “lifespan” as a keyword in Pubmed; the transcript/protein of the gene correlates with average lifespan; the gene/protein has an expression *cis*-eQTL or a protein *cis*-pQTL; the RNAi clone with the right sequence is available from at least one of the libraries (Ahringer and Vidal libraries). Grey color: not available (mRNA or protein not measured). Top candidate genes, *rict-1* and *gfm-1* are highlighted in red and bold. *: known longevity gene. **Related to Figures S7, S8 and Table S5.**

We further explored other life-history traits and detected four significant QTLs: one for progeny emergence (the time when the 1^st^ progeny is observed) on Chr. V:12,125,475, one for egg emergence (the time when the 1^st^ egg is observed, indicating sexual maturity) on Chr. V:20,279,818, and two for shapes straight and supercoil, both on Chr. X (X: 11,549,662 and X:12,745,016 respectively) (Figure 5A and Table S5). When examining the locus at V:12,125,475, we found that RIAIL strains with the N2 genotype exhibited a longer time for progeny emergence compared to those with the CB4856 genotype (Figure 5C). Furthermore, despite observing a correlation between developmental time and lifespan traits (Figures 2F-2G), we did not detect any significant or shared QTL between the two traits, suggesting that this correlation does not necessarily imply a common genetic regulation.

### Exploration of lifespan QTL identified *gfm-1* as the top candidate gene

To identify potential mediators of the effect of the lifespan QTL, we performed differential expression analysis between RIAIL strains with the CB4856 genotype at that locus compared to those with the N2 genotype. Although only a limited number of genes were differential expressed (adjusted p-value < 0.05, Table S5), GSEA revealed significant differences at the pathway level (Figure 5D). We noted a greater number of significantly positively enriched genesets related to cellular components compared to biological processes (Figure 5D). Neuropeptide signaling and G protein-coupled receptor signaling pathways were significantly up-regulated in strains carrying the CB4856 marker, which exhibited an extended lifespan. The cellular component results highlight a significant upregulation of genes involved in organellar/mitochondrial ribosome-associated with the genetic variation at this lifespan locus (Figure 5D).

The lifespan QTL encompassed eight genes. To prioritize the most likely candidate modulators of lifespan, we considered a wide range of factors (Figure 5E), namely whether there were any genetic variants in the gene between N2 and CB4856, whether any of these were mis-/nonsense mutations, the presence of *cis-*e/pQTLs defined as genomic loci near the gene of interest (in *cis*) that explain the variation in expression levels of mRNA (eQTL) or protein (pQTL) in of that gene, whether the gene was differentially expressed between strains with N2 vs. CB4856 genotypes, prior knowledge of the gene being associated with aging (in GenAge, a curated database of genes associated with age-related processes, (Tacutu et al., 2018)) or lifespan (PubMed searching for “lifespan”), and whether the gene was correlated with lifespan at the mRNA or protein level. Most of the genes had some genetic variants, many with missense or nonsense variants as well, but among these, only *rict-1* met additional criteria as it has been reported to be associated with both aging and lifespan. *rict-1* encodes a key component of the mTORC2 complex and loss-of-function mutations have previously been shown to increase the lifespan of *C. elegans* in specific conditions (Mizunuma et al., 2014; Soukas et al., 2009). In addition, we were interested in *gfm-1,* as mitochondria play a key role in longevity regulation and *gfm-1* is a known mitochondrial gene, encoding the G elongation factor mitochondrial 1.

### RNAi of *gfm-1* prolonged lifespan by activating the UPR^mt^

To examine the causal relationship between these candidate genes unveiled hitherto and longevity modulation, we knocked down the candidate genes by feeding worms with RNAi bacteria targeting each candidate gene, starting from the maternal phase, and measured their lifespans (Figure 6A). Knockdown of *gfm-1* showed the most significant lifespan extension compared to the other candidate genes (p-value < 0.0001) (Figure 6A). Besides *gfm-1*, our survival analysis revealed that knocking down of *mltn-1* (molting cycle MLT-10-like protein, (Meli et al., 2010)) also significantly prolonged lifespan (p-value < 0.001), albeit to a lower extent. Conversely, the knockdown of *rict-1* resulted in a shorter lifespan. This is potentially due to the adverse effects of RNAi treatment from an early developmental stage as previous studies exposed worms to *rict-1* RNAi bacteria exclusively during adulthood to bypass the developmental functions of TORC2 and observed a prolonged lifespan (Mizunuma et al., 2014; Robida-Stubbs et al., 2012).

**Figure 6.**
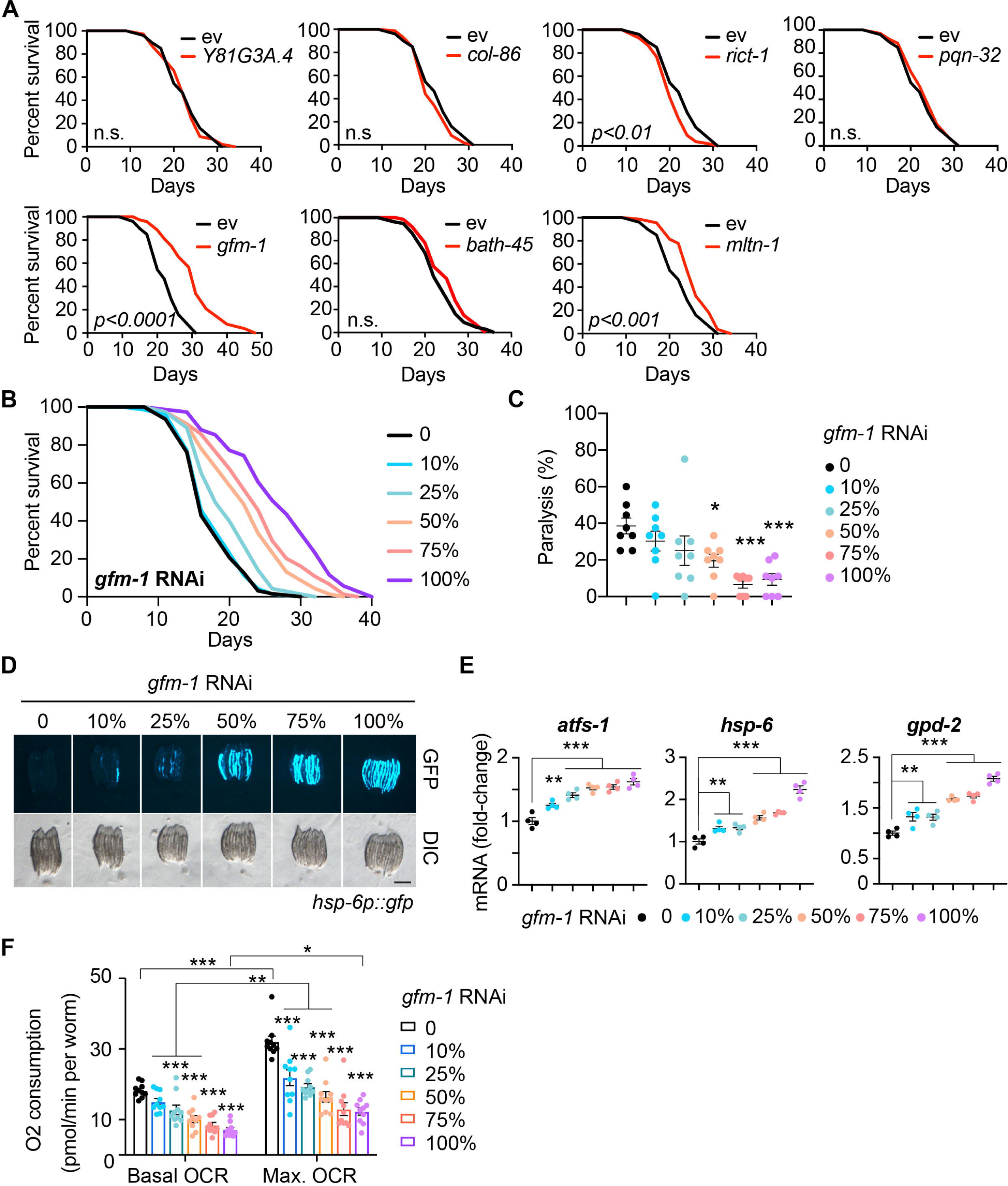
RNAi of *gfm-1* induced UPR^mt^ activation and prolonged lifespan in *C. elegans*. (**A**) Lifespan of worms fed with ev (control RNAi) (black) or candidate gene RNAi (red). P-values represent a comparison with the controls calculated using the log-rank test. n.s.: not significant. (**B**) RNAi of *gfm-1* extends worm lifespan in an RNAi dose-dependent manner. Worms fed with ev (control RNAi) or 10%-100% *gfm-1* RNAi; control RNAi was used to supply to a final 100% of RNAi. (**C**) Age-related paralysis of worms fed with ev or 10%-100% *gfm-1* RNAi. Error bars denote SEM. Statistical analysis was performed by one-way ANOVA followed by Tukey post-hoc test (*p < 0.05; ***p < 0.001). (**D**) RNAi of *gfm-1* induced the UPR^mt^ (*hsp-6p::gfp* reporter) in a dose-dependent manner. Worms fed with ev or 10%-100% *gfm-1* RNAi. Scale bar: 0.5 µm. (**E**) RT-qPCR results of mRNA levels (n = 4 biological replicates) in worms fed with ev, or 10%-100% *gfm-1* RNAi. Statistical analysis of RT-qPCR results was performed by one-way ANOVA followed by Tukey post-hoc test (**p < 0.01; ***p < 0.001). Values in the figure are mean ± SEM. (**F**) RNAi of *gfm-1* reduced both basal and max. oxygen consumption rate (OCR) compared to those of controls on day 1 of adulthood. Values in the figure represent mean ± SEM. Statistical analysis of RT-qPCR results was performed by one-way ANOVA followed by Tukey post-hoc test (*p < 0.05; **p < 0.01; ***p < 0.001). **Related to Figure S9.**

To further characterize the mechanism of *gfm-1* RNAi*-*mediated longevity, we conducted several functional assays. We assessed the effect of *gfm-1* RNAi on lifespan and healthspan with a dilution of RNAi bacteria, including 10%, 25%, 50%, 75%, and 100% (control RNAi was used to supply to a final 100% of RNAi for all conditions). Worms exposed to different amounts of *gfm-1* RNAi showed a dose-dependent lifespan extension (Figure 6B) and reduction of age-related paralysis (Figure 6C). Because *gfm-1* encodes a mitochondrial translation elongation factor, we considered whether the mitochondrial stress response (MSR), through components such as the mitochondrial unfolded protein response (UPR^mt^), was involved in longevity changes observed with *gfm-1* knockdown. Indeed, *gfm-1* RNAi robustly increased the GFP expression of *hsp-6p::gfp* worms and significantly upregulated the expression of the UPR^mt^ genes, including *atfs-1*, *hsp-6*, and *gpd-2* (Figures 6D-6E). In line with this, mitochondrial respiration was also reduced upon *gfm-1* knock down in a dose-dependent manner (Figure 6F). These results confirmed the beneficial effect of mitochondrial inhibition and UPR^mt^ activation on healthy aging and longevity (Durieux et al., 2011; Houtkooper et al., 2013).

Furthermore, we investigated the potential mechanism of *mltn-1* RNAi-induced longevity by examining whether any of the established longevity pathways contribute to the observed lifespan extension (Figure S9). We fed *mltn-1* RNAi to worms with mutations mimicking caloric restriction (*eat-2* mutant and *sir-2.1* overexpression worms) (Lakowski and Hekimi, 1998; Tissenbaum and Guarente, 2001), insulin/IGF-1 signaling (*daf-2* mutant) (Kenyon et al., 1993), AMPK signaling (*aak-2* mutant) (Schulz et al., 2007) and oxidative stress response (*skn-1* mutant) (Lehrbach and Ruvkun, 2016) (Figure S9). Of note, *mltn-1* RNAi prolonged the lifespan of worms overexpressing *sir-2.1* overexpression and *skn-1* mutants, indicating that *mltn-1* RNAi regulates longevity independent of sirtuin-induced caloric restriction and oxidative stress response (Figures S9B and S9E). However, *mltn-1* RNAi did not further extend the lifespan of *eat-2* and *daf-2* mutants (Figures S9B and S9C), and the lifespan extension induced by *mltn-1* RNAi was almost completely abolished in *aak-2* mutant worms (Figure S9D). These results suggest that the knockdown of *mltn-1* extends worm lifespan in an AMPK-dependent manner and potentially mimics caloric restriction. Taken together, these findings further reinforced the assertion that our approach enabled us to identify novel inducers of longevity.

### Variants in human *GFM1* and *RICTOR* elevated risk of cardiovascular conditions, dementia, diabetes, kidney, liver diseases, and death

Age-related diseases play a significant role in shaping longevity (Franceschi et al., 2018). To explore the human relevance of our newly identified longevity genes, we took advantage of the UK Biobank, a large-scale population-based cohort study with extensive health and medical information (Bycroft et al., 2018). While *mltn-1* is a *C. elegans-*specific gene, we identified *GFM1* and *RICTOR* as the human orthologs of worm *gfm-1* and *rict-1*, and then used Cox proportional-hazards models (Borgan, 2001; Cox, 1972; Therneau, 2023) to investigate whether variants within these genes were associated with disease risk (Figure 7A). In addition, given that *rict-1* is the gene encoding the mTORC2 component Rictor, we also investigated the potential association of human *RICTOR* with diseases in the UK Biobank (Figure 7A). We explored the association between single nucleotide polymorphisms (SNPs) in *GFM1* and *RICTOR* (selection based on criteria outlined in the STAR method) and the lifelong incidence of 19 diseases as well as all-cause mortality (referred to as “Death”) (Tables S6 and S7). 21 GFM1 SNPs showed an association with 13 diseases and death (Benjamini-Hochberg adjusted p-value < 0.05), and 59 RICTOR SNPs correlated with 18 diseases and death. All associations showed an increased risk for diseases with the alternate (minor) allele. The majority of *GFM1* SNPs were linked to myocardial infarction, cardiomyopathy, heart failure, Alzheimer’s disease/dementia, diabetes, kidney failure, liver disease, and death. Similarly, *RICTOR* SNPs were associated with these diseases as well as cerebrovascular disease, vascular dementia, and Parkinson’s disease (Figure 7B). Taken together, the identification of these SNPs of *GFM1* and *RICTOR* offers new insights into the genetic underpinnings of many age-related disease diseases. Specifically, our results highlight distinct genomic regions implicated in the predisposition to cardiomyopathy and heart failure for *GFM1*, and heart failure and aortic aneurysm for *RICTOR*, diseases that negatively affect survival in humans.

**Figure 7.**
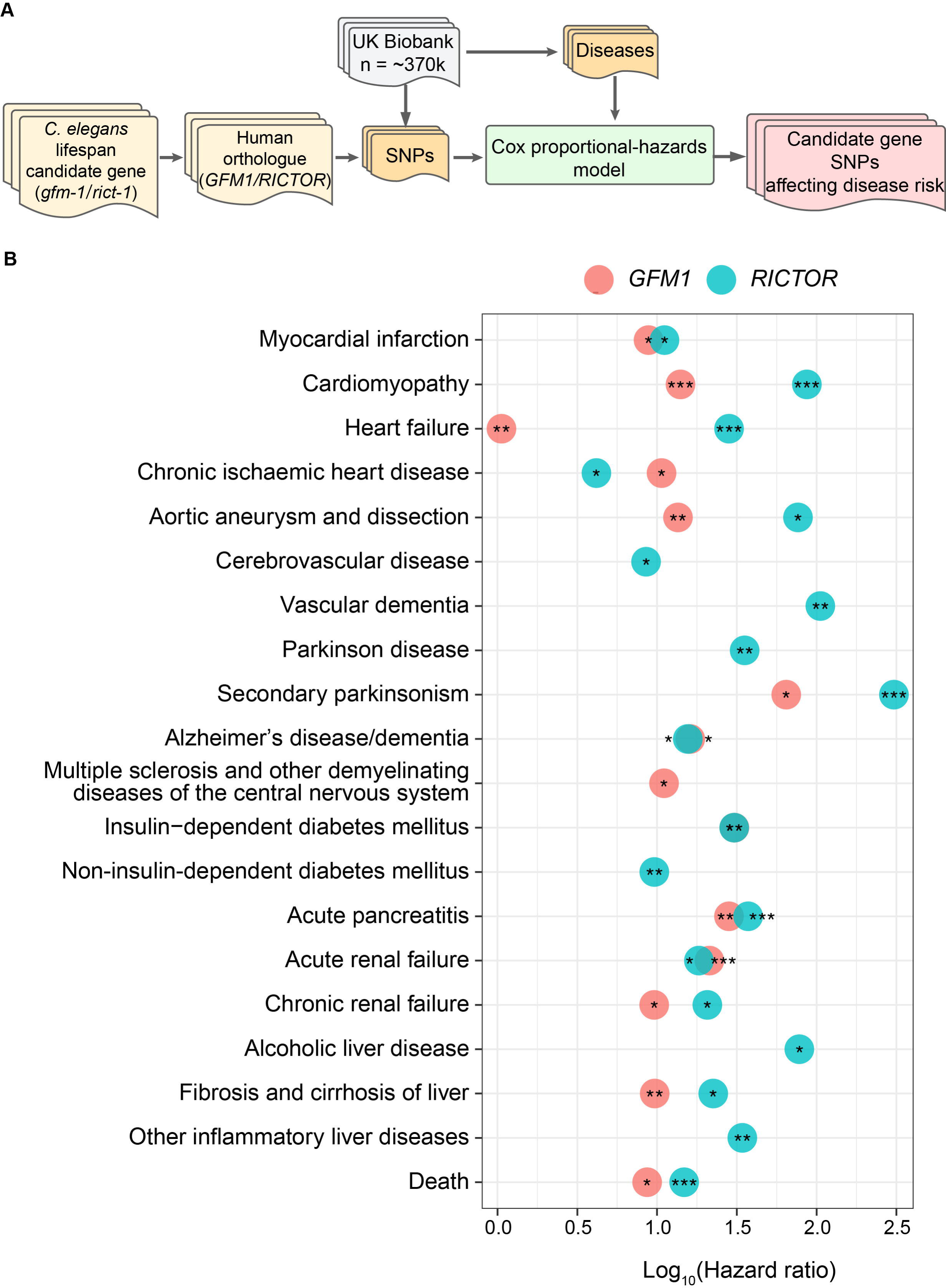
Exploration of human *GFM1* and *RICTOR* SNP association with disease incidence in the UK Biobank. (**A**) The workflow of disease risk associations between *GFM1*/*RICTOR* variants and life-long incidence of diseases and all-cause mortality with Cox proportional-hazard models. (**B**) Identification of 21 and 59 SNP-disease associations (BH-Adjusted p value < 0.05) for *GFM1* (colored by light red) and *RICTOR* (colored by light blue). 19 diseases (including cardiovascular diseases, dementia, diabetes, renal failure, and liver disease) and death are pre-selected. The variant-disease associations for each disease are ranked by BH-adjusted p-value, and only the most significant one is shown in the plot. ***: BH-adjusted p-value < 0.001; **: BH-adjusted p-value < 0.01; *: BH-adjusted p-value < 0.05. The Hazard ratio calculated by Coxproportional-hazard models is indicated in the X-axis. **Related to Tables S6 and S7.**

## Discussion

Here we present a multi-omics atlas of the worm RIAILs, as a resource to understand the regulation of longevity. The observed difference in average lifespan between the parental strains was consistent with previous studies (Gao et al., 2022; Lee et al., 2016). The RIAIL strains exhibited extensive lifespan variation with some strains exceeding that of the parentals suggesting the presence of transgressive segregation (Rieseberg et al., 1999). Research across species has revealed consistent trade-offs that influence lifespan and life-history traits, with correlations observed between key phenotypic traits such as lifespan and fecundity (Luckinbill et al., 1984), development time and fecundity (Ghalambor et al., 2004), development time and lifespan (Marchionni et al., 2020), as well as body size and longevity (Bou Sleiman et al., 2022; de Magalhaes et al., 2009). However, research exploring the correlations between longevity and early life history traits in wild *C. elegans* populations is relatively scarce (Anderson et al., 2011; Lee et al., 2016). We therefore tracked various life-history phenotypes during the early life stage of the RIAILs, gathering data on developmental progression, reproductive capability, fertility, and behavioral activity. Only developmental time and egg emergence – both reflecting sexual maturity – had a weak to moderate correlation with lifespan, which is in line with the absence of any overlapping QTL between early life history and lifespan traits. These data are corroborated by a prior study in worms, which proposed that development, reproduction, and lifespan are under independent genetic regulation (Johnson, 1987), and work in *D. melanogaster*, where a disconnect between life-history traits and lifespan was observed when examining variations in larval food conditions (Tu and Tatar, 2003; Zwaan et al., 1992). In a similar vein, fly selection experiments have yielded inconsistent results in terms of discovering genetic correlations between development time, body size, and longevity (Chippindale et al., 1994; Hoedjes et al., 2019; Zwaan et al., 1995a, b).

Subsequently, we investigated whether multi-omic molecular characteristics, such as gene expression, protein, or lipid abundance, could be linked to lifespan in the RIAILs. We did not detect any significant correlations between individual transcript/protein/lipid and lifespan traits following multiple testing corrections possibly due to small marginal effects or more complex gene interactions. These correlations, however, strengthened and reached statistical significance when we performed the gene set enrichment analysis on genes ranked by the mRNA-lifespan associations, supporting the presence of numerous biological pathways that are potentially involved in the modulation of RIAIL lifespans. The lack of significant associations at the individual transcript level may not negate the possibility of a functional impact and physiological relevance at the pathway level, where complex interactions and synergistic effects may come into play. For instance, neuropeptide signaling and G protein-coupled receptor (GPCR) signaling were particularly notable among the pathways that were positively associated with lifespan traits. This finding aligns with prior studies, where one demonstrated the role of the neuropeptide signaling pathway in extending *C. elegans* lifespan (Frakes et al., 2020; Savini et al., 2022), and another highlighted the influence of the GPCR pathway on longevity across humans and various animal models including worms (Lagunas-Rangel, 2022). Moreover, when we examined the genetic determinants of lifespan by QTL mapping, neuropeptide signaling and GPCR pathways were also upregulated dependent on the genotype at the identified lifespan locus on Chromosome II. This consistent pattern suggests that the association between the transcriptome and lifespan was influenced by this specific lifespan locus. While we identified intriguing gene sets associated with lifespan traits at the transcript level, these associations were not replicated in the analysis between protein expression and lifespan traits. The discrepancy between the transcriptomic and proteomic levels could be attributed to several factors, such as post-transcriptional regulation, protein turnover, limitations in the proteomic detection methods (Schubert et al., 2017), or differential effect of natural variation on the proteome (Kamkina et al., 2016). These discrepancies emphasize the importance of considering multiple omics layers to obtain a comprehensive understanding of biological processes and their role in determining phenotypic outcomes.

Alterations in circulating lipid concentrations, triggered by genetic influences, lifestyle choices, and environmental conditions, can escalate the risk of age-associated disorders (Harshfield et al., 2021). We therefore also collected full lipidomic profiles of RIAILs and investigated whether complex lipids might also be associated with specific lifespan traits. We found that TGs, CLs, and PIs were over-represented in positive lipid-lifespan associations, while PEs were enriched in negative lipid-lifespan associations. It is notable that CLs, comprised mainly of polyunsaturated fatty acid chains, were found to be among those positively associated with lifespan traits. CLs are mitochondria-specific phospholipids essential for preserving mitochondrial integrity (Falabella et al., 2021). Due to their special cellular confinement, CLs are closely related to the maintenance of mitochondrial function, which connects CLs to longevity and the progression of age-related disease (Dai et al., 2021). This aligns with our findings that indicate a positive correlation between CLs and various lifespan traits. In contrast, the level of PEs consisting of less saturated fatty acids exhibited a negative correlation with lifespan traits. Although PEs are the second most prevalent glycerophospholipid in eukaryotic cells and positively regulate autophagy and lifespan in yeast and mammalian cells (Calzada et al., 2016; Rockenfeller et al., 2015), decreased levels of PE were associated with lower beta-amyloid accumulation in both mammalian cells and flies (Nesic et al., 2012), suggesting a complex role of PEs in regulating age-related effects and longevity.

In addition to the multi-omic characterization of the RIAIL population, we performed QTL mapping and identified candidate lifespan loci on Chr. II, with strains carrying the CB4856 genotype showing longer lifespans compared to those with the N2 genotype at this locus. This finding was particularly interesting considering that the N2 parental strain displayed a significantly longer lifespan compared to the CB4856 strain. RNAi against the seven candidate genes in that locus found that knockdown of *gfm-1* and *mltn-1* led to significant lifespan extensions. Our analyses suggest that the dose-dependent lifespan extension and reduction of age-related paralysis through *gfm-1* inhibition could be mediated by the modulation of the mitochondrial stress response. Although we did not detect an mRNA or protein QTL for *gfm-1* within the same lifespan locus, the experimental findings were in line with this gene encoding the G elongation factor mitochondrial 1 and the upregulation of gene sets associated with mitochondrial ribosomes at this locus in the RIAILs population. Given that *mltn-1* is specific to *C. elegans*, its translational relevance to human studies is uncertain. Another candidate that is known as a longevity gene is *rict-1*. As an essential component of the TORC2 complex, *rict-1* (Rictor) is vital for development, which likely explains why worms subjected to *rict-1* RNAi have shortened lifespans. Several studies, however, have demonstrated an increased lifespan in worms fed with *rict-1* RNAi when TORC2 activity is attenuated specifically during adulthood (Mizunuma et al., 2014; Robida-Stubbs et al., 2012). To evaluate the potential clinical relevance of the selected candidate genes that are conserved in humans, we took advantage of the UK biobank and demonstrated that variants in the human *GFM1* and *RICTOR* genes were associated with several age-related and metabolic diseases, including heart disease, dementia, diabetes, renal failure, liver disease, all of which contribute to shorter life expectancy due to their detrimental effects on cardiac, metabolic and overall health and organ function.

In summary, our study unveiled a specific genetic locus that plays a role in determining lifespan variation within the RIAIL population. Furthermore, we identified known and novel longevity modulators, including *rict-1*, *gfm-1*, and *mltn-1*, which we validated experimentally. The comprehensive multi-layered characterization of the RIAIL population is now also made accessible through an open-access web resource (https://lisp-lms.shinyapps.io/RIAILs/), which provides a valuable tool for investigating the intricate relationships between biochemical and whole-body phenotypes and for hypothesis generation for the scientific community.

### Limitations of the study

We note several limitations and future directions of our work. First, the relatively low sample size of worms (60 worms/strain) used for lifespan analysis restricts our ability to get an accurate estimate of late-life mortality, especially for the maximal lifespan of the strain (Brooks et al., 1994; Carey et al., 1992). This likely undermines our statistical power in evaluating the associations between traits and late-lifespan phenotypes. Second, the life-history trait screening was done in liquid culture using the microfluidics device, while the lifespan assays were performed on plates; we can hence not exclude a possible influence of different culture conditions on traits. Third, the gathered molecular characteristics encompass aggregated data at the strain level and are limited to a single early time point. However, expanding the data collection to include later time points would enable the exploration of age-related dynamics associated with these traits. Finally, the experimental validations of *gfm-1*, *rict-1*, and *mltn-1* were conducted using RNAi knockdown in the N2 Bristol background. Moving forward, an important avenue for further investigation would involve utilizing CRISPR technology to examine the specific variant of *gfm-1* in the RIAILs population.

## Acknowledgments

We thank Andersen’s lab and *Caenorhabditis* Genetics Center for providing the *C. elegans* strains. We thank all members of Johan Auwerx and Kristina Schoonjans laboratories for helpful discussions. We also thank Dr. Changliang Chen from the Department of Medicine, University of Wisconsin Carbone Cancer Center, University of Wisconsin-Madison, School of Medicine and Public Health, for help in the LS-MS sample preparation. This work was supported by grants from the EPFL, the European Research Council (ERC-AdG-787702), the Swiss National Science Foundation (SNSF 31003A_179435 and Sinergia CRSII5_202302), and GRL grant of the National Research Foundation of Korea (NRF 2017K1A1A2013124). The work in the lab of J.J.C. was supported by P41GM108538 (J.J.C.) and R35GM118110 (J.J.C.) from the National Institutes of Health. A.W.G. was supported by the United Mitochondrial Disease Foundation (PF-19-0232), Amsterdam UMC Postdoc Career Bridging Grant (2021), Horizon-MSCA-PF-EF-2022 (101108082), and AGEM Talent Development Grant (2023). T.Y.L. was supported by the “Human Frontier Science Program” (LT000731/2018-L).

## Author contributions

A.W.G. and J.A. conceived the project. A.W.G. performed all the lifespan measurements of RIAILs. E.K. collected early life-history phenotypes of RIAILs using a Nagi Bioscience’s high-content screening device. A.W.G. and T.Y.L. collected all the worm samples for multi-omics measurements. Y.Z. measured proteomics and lipidomics profiles. G.E.A., J.D.M., J.S., X.L., and A.W.G. performed data analysis, with great help from Y.Z., E.K., K.A.O., A.L., L.M., M.C., and M.B.S. for confirmation. A.W.G. and W.L. performed the validation experiments. J.A., J.D.M., R.H.H., and J.J.C. supervised the study. A.W.G., G.E.A., and J.A. wrote the manuscript with comments from all the co-authors. This research has been conducted using the UK Biobank Resource under Application Number 48020.

## Competing interests

J.J.C. is a consultant for Thermo Scientific, Seer, and 908 Devices. E.K., L.M., and M.C. are employees of Nagi Bioscience S.A. Other authors do not declare a conflict related to this study.

**Correspondence and requests for materials** should be addressed to J.A.

## STAR METHODS

- KEY RESOURCE TABLE
- RESOURCE AVAILABILITY

- Lead contact
- Data and code availability
- EXPERIMENTAL MODEL AND SUBJECT DETAILS

- *C. elegans* and bacterial feeding strains
- METHOD DETAILS

- Lifespan measurements
- Phenotyping by microfluidics
- Sample collection for RNA-seq, proteomics, and lipidomics analysis
- RNA extraction and RNA-seq analysis
- Protein digestion
- Lipid extraction
- LC-MS setup
- Data analysis for proteomics and lipidomics
- Survival analysis and lifespan traits extraction
- Life-history trait batch correction
- Statistical analysis
- Dirichlet regression analysis
- Variant calling and genetic map construction
- Association mapping and gene-set enrichment analysis
- Lipid class over-representation analysis
- Quantitative trait locus (QTL) mapping
- Lifespan locus differential analysis and gene-set enrichment analysis
- UK Biobank *GFM1* SNP-disease time-to-event analysis
- Figures and visualizations
- Data availability
- Fluorescent image for assessing the UPR^mt^ activation
- Real-time quantitative PCR (RT-qPCR)
- OCR measurements by Seahorse
- QUANTIFICATION AND STATISTICAL ANALYSIS

## RESOURCE AVAILABILITY

### Lead contact

Further information and requests for resource and reagents should be directed to and will be fulfilled by the Lead Contact, Johan Auwerx.

### Data and code availability

- The RNA-seq data has been deposited in the National Center for Biotechnology Information Gene Expression Omnibus database (accession number: GSE252593). The Mass spectrometry raw files have been deposited to the MassIVE database (accession number MSV000088622; ftp://MSV000088622@massive.ucsd.edu).
- This study did not generate any codes.
- Any additional data types/resources will be shared by the lead contact upon request after publication.

### Key Resources Table

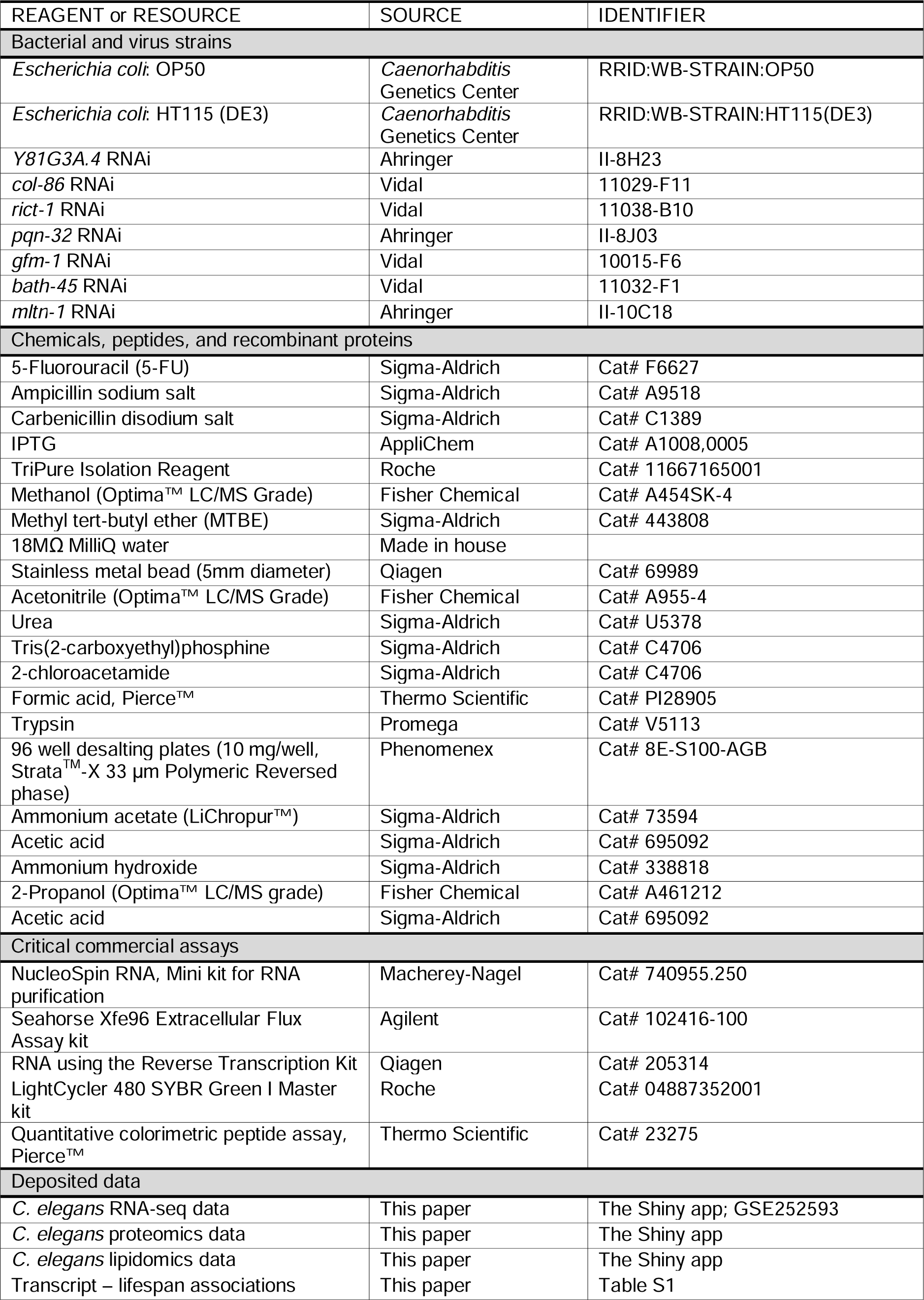
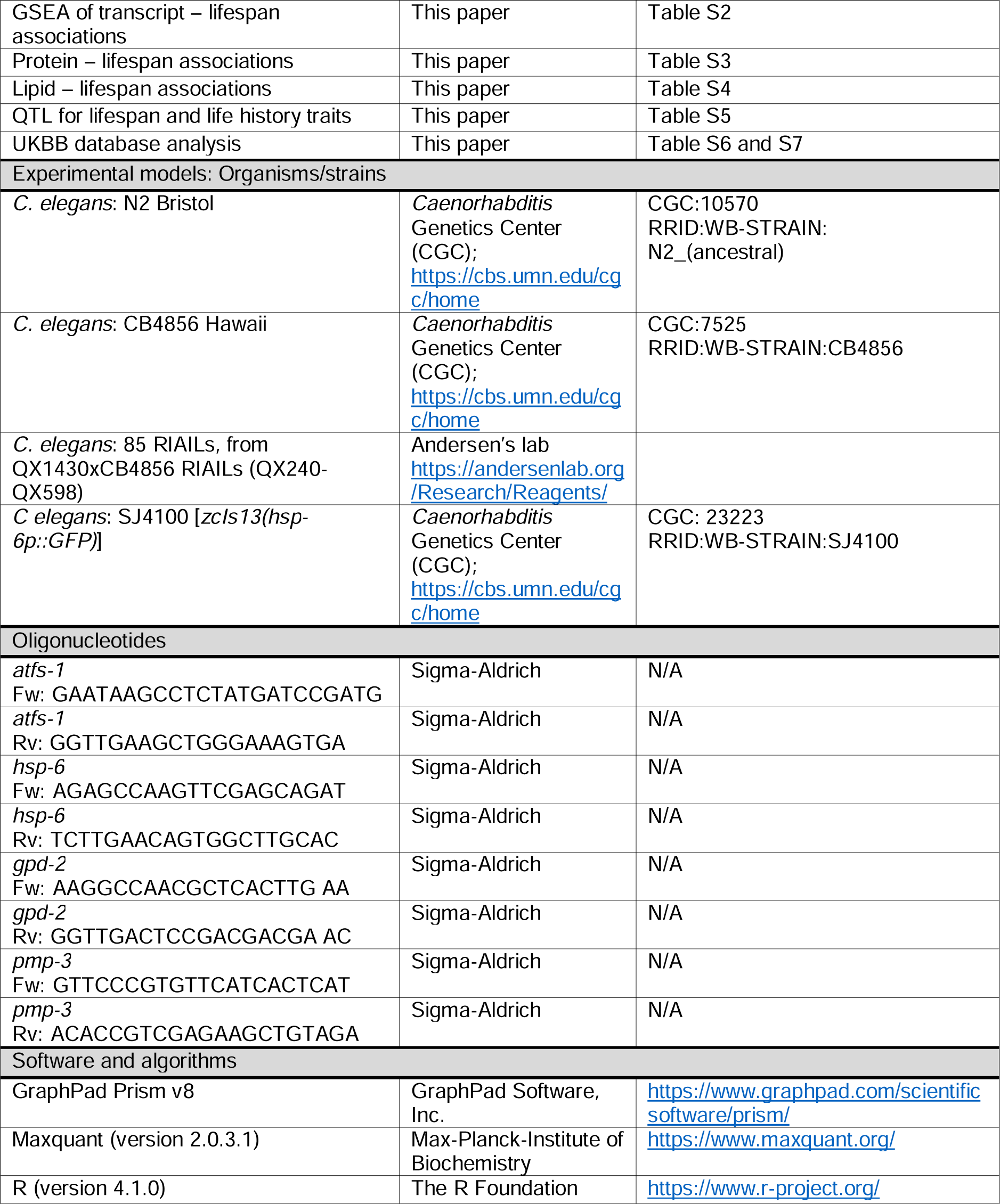
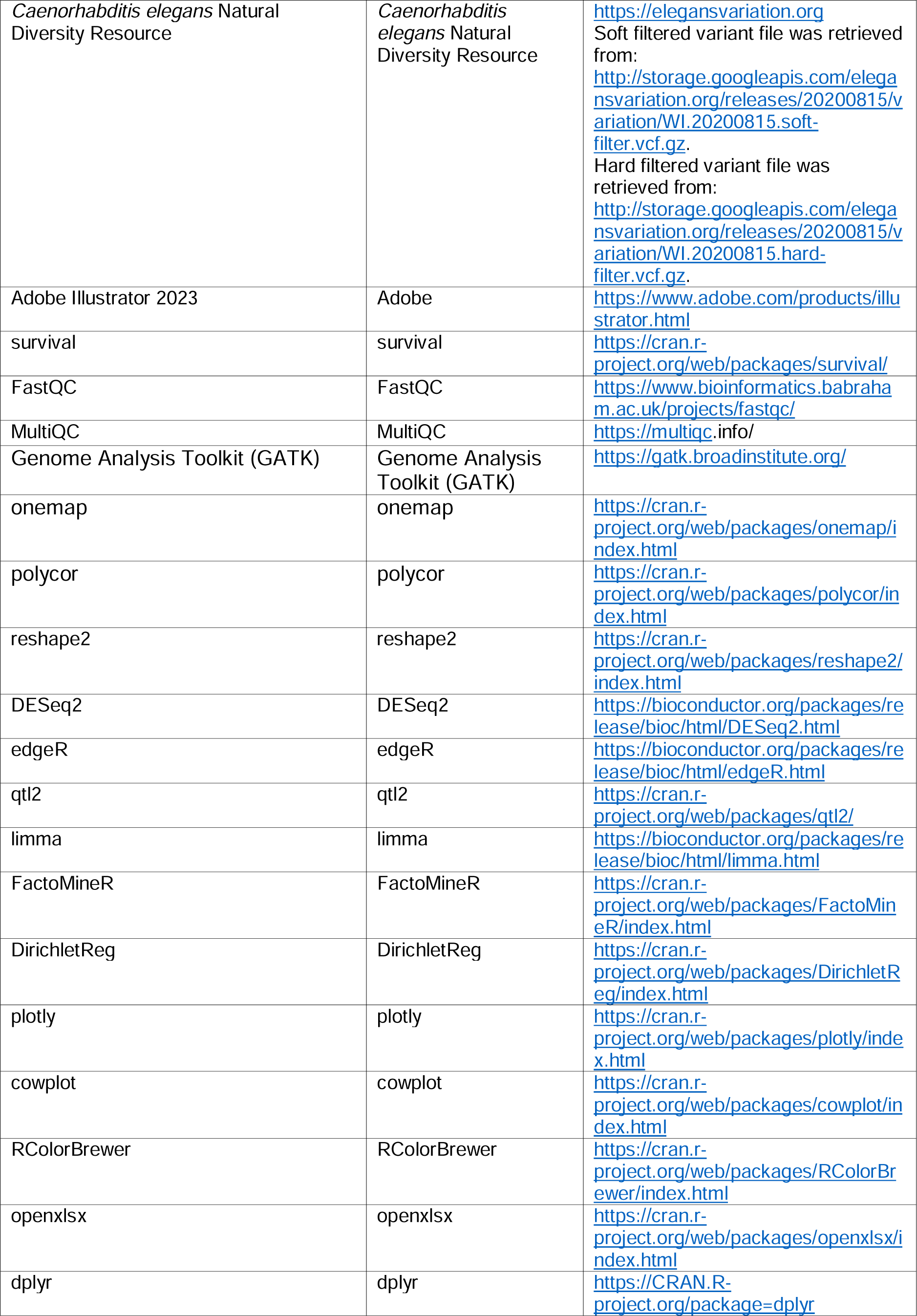
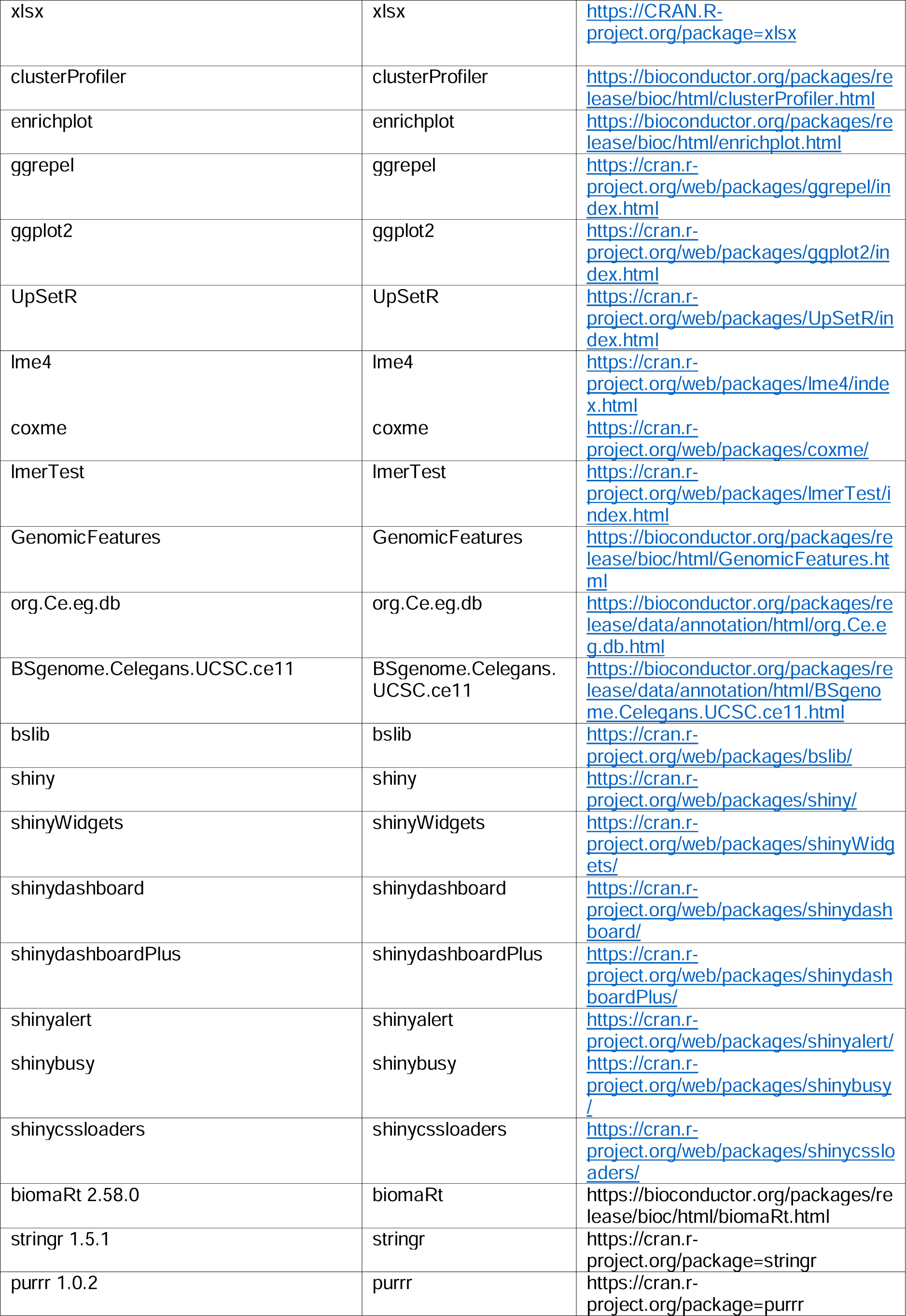
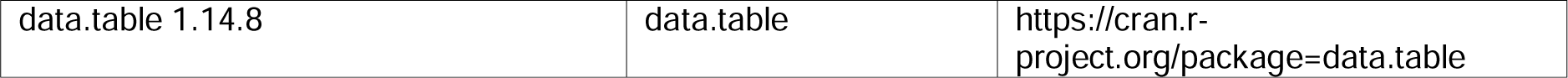

## EXPERIMENTAL MODEL AND SUBJECT DETAILS

### Bacterial strains and C. elegans strains

The Bristol strain (N2) and Hawaii strain (CB4856) were used as the wild-type strains, and SJ4100 *[zcIs13(hsp-6p::GFP)]*, DA465 *[eat-2(ad465)]*, *[sir-2.1(ok434)]*, CB1370 *[daf-2(e1370)]*, RB754 *[aak-2(ok524)]*, and GR2245 *[skn-1(mg570)]* were obtained from the *Caenorhabditis* Genetics Center (CGC; Minneapolis, MN). *E. coli* OP50 and HT115 strains were also obtained from the CGC. RNAi clones against *Y81G3A.4*, *col-86*, *rict-1*, *pqn-32*, *gfm-1*, *bath-45*, and *mltn-1*, were obtained from the Ahringer and Vidal libraries and verified by sequencing before use (detailed in the Key Resource Table). Worms were cultured and maintained at 20°C and fed with *E. coli* OP50 on Nematode Growth Media (NGM) plates unless otherwise indicated.

## METHOD DETAILS

### Lifespan and paralysis measurements

Lifespan was measured as described (Mouchiroud et al., 2011). In general, 5-10 L4 worms of each worm strain were transferred onto RNAi plates (containing 2 mM IPTG and 25 mg/mL carbenicillin) seeded with *E. coli* HT115 bacteria or RNAi bacteria. After the F1 progenies reached the last larval stage L4, worms were then transferred onto RNAi plates containing 10 µM 5FU. Approximately 60 worms were used for each condition and scored every other day. For the validation experiments of the candidate genes, 80 worms were used for each condition.

Paralysis was manually assessed through the previously described poking method (McColl et al., 2012), with a minimum of 80 worms analyzed per condition.

### Phenotyping by microfluidics

The phenotypic readouts reflecting development, growth dynamics, fertility, and reproduction of different RIAIL strains were collected with the SydLab robotic microfluidic-based platform developed by Nagi Bioscience SA, which allows high-throughput and high-content *C. elegans* screenings. A synchronized population of *C. elegans* was injected into microfluidic chips at the L1 larval stage. Worms were confined within dedicated microfluidic chambers and were continuously fed with freeze-dried *E. coli* OP50 solution. The images of each chamber were recorded every hour for the whole duration of the experiment. At the end of the experiment the collected images were processed by a set of software modules (also developed by Nagi Bioscience) based on machine learning algorithms, allowing a fully automated and standardized way for feature extraction and data analysis. The experiments were performed at 23°C.

### Sample collection for RNA-seq, proteomics, and lipidomics analyses

Worms of each RIAIL strain were cultured on plates seeded with *E. coli* OP50, and then worm eggs were obtained by alkaline hypochlorite treatment of gravid adults. A synchronized L1 population was obtained by culturing the egg suspension in sterile M9 butter overnight at room temperature. Approximately 2000 L1 worms of each RIAIL strain were transferred onto plates seeded with *E. coli* HT115. L4 worms were harvested after 2.5 days with M9 buffer and washed three times. Worm pellets were immediately submerged in liquid nitrogen for snap-freezing and stored at −80°C until use.

### RNA extraction and RNA-seq data analysis

On the day of the RNA extraction, 1 mL of TriPure Isolation Reagent was added to each sample tube. The samples were then frozen and thawed quickly eight times with liquid nitrogen and a 37 °C water bath to rupture worm cell membranes. Total worm RNA was extracted by using a column-based kit from Macherey-Nagel. RNA-seq was performed by BGI with the BGISEQ-500 platform. FastQC (version 0.11.9) was used to verify the quality of the reads (de Sena Brandine and Smith, 2019). Low-quality reads were removed and no trimming was needed. Alignment was performed against worm genome (WBcel235 sm-toplevel) following the STAR (version 2.73a) manual guidelines (Dobin et al., 2013). The STAR gene-counts for each alignment were analyzed for differentially expressed genes using the R package DESeq2 (version1.32.0)(Love et al., 2014) using a generalized-linear model. Count data were normalized to counts per million (CPM) using edgeR (version 3.36.0) for visualizations of expression data. Biological process (BP) overrepresentation analysis and Gene Set Enrichment Analysis (GSEA) were performed using Clusterprofiler (version 4.2.2) and org.Ce.eg.db (version 3.14.0). A principal component analysis was also generated to explore the primary variation in the data (Lê et al., 2008; Risso et al., 2014). For RT-qPCR, worms were collected and total RNA was extracted as described for the RNA-seq sample preparation. cDNA synthesis was conducted from total RNA by the Reverse Transcription Kit (Qiagen, Cat# 205314). qPCR was performed using the Light Cycler 480 SYBR green I Master kit (Roche, Cat# 04887352001). The primers used for RT-qPCR are listed in the Key Resource Table. *pmp-3* was used as housekeeping controls.

### Lipid extraction

The extraction procedures have been described previously (Zhu et al., 2023). All reagents were chilled on ice and samples were maintained at ≤ 4°C during the extraction procedure. A metal bead was added to each sample. Next, 500 µL M1 (tert-Butyl methyl ether:Methanol = 3:1, v:v) was added to each tube and vortexed for 2 minutes. 325 µL M2 (H_2_O: Methanol = 3:1, v:v) was added to each tube. Samples were vortexed briefly. Then, samples were flash-freezed in liquid nitrogen and thawed on ice. This step was done three times to facilitate cell breakage. Samples were transferred to a bead-beater and shaken at 1/25 s frequency for 5 min, and this process was done three times. The samples were then centrifuged for 10 min at 12,500 g at 4°C. For downstream lipid analysis, 200 µL of the organic layer (upper phase) was transferred to a glass autosampler vial and dried by vacuum centrifugation. Remaining protein pellets on the bottom were kept on ice until further digestion.

Once dried, organic extracts intended for lipid analysis were resuspended in 100 µL 65:30:5 Isopropanol:Acetonitrile:Water and vortexed for 20 s prior to analysis by Liquid chromatography–mass spectrometry (LC-MS). Aqueous extracts intended for metabolomic analysis were resuspended in 50 µL 1:1 Acetonitrile (ACN):Water and also vortexed for 20 s prior to analysis by LC-MS.

### Protein digestion

Remaining protein pellets on the bottom were washed with 1 mL ACN and centrifuged at 10 kg for 3 min at 4 °C. Supernatant ACN was aspirated and the protein pellets sit for 10-15 min at room temperature, or vacuum dried briefly to dry up the liquid in the bottom of the tube. 300 μL lysis buffer (8M urea with 100 mM tris(2-carboxyethyl)phosphine, 40 mM chloroacetamide and 100 mM tris (pH = 8.0) was added to each sample and vortex till the protein pellets were fully dissolved. 5 μg LysC was added to each sample with protein:enzyme ratio 70:1 (digestion lasted overnight at room temperature). Trypsin at 70:1 protein:enzyme was added to each sample after diluting the lysis buffer to 2 M urea and digestion lasted for six hours at room temperature. Desalting was carried out with 96 well desalting plates. A blank well between any two samples was reserved to avoid cross contamination. Desalting started with equilibrating the desalting wells with 1 mL 100% ACN, followed by 1 mL 0.2% FA. Acidified peptide mixture was loaded to the 96 well desalting plate, followed by 2 mL 0.2% FA wash. Peptides were eluted into a 96-well collection plate with 600 μL 80% ACN with 0.2% FA. Peptides were vacuum dried down and stored in −80°C freezer until resuspension with 0.2% FA. After resuspension, peptide concentration was measured using a quantitative colorimetric peptide assay.

### LC-MS setup

#### Proteomics

Peptides were separated on an in-house prepared high pressure reversed phase C18 column (Shishkova et al., 2018). Briefly, a 75-360 μm inner-outer diameter bare-fused silica capillary was packed with 1.7 μm diameter, 130 Å pore size, Bridged Ethylene Hybrid C18 particles (Waters) under high pressure of 25K psi to a final length of ∼40 cm. The column was installed onto a Thermo Ultimate 3000 nano LC and heated to 50 °C for all runs. Mobile phase buffer A was composed of water with 0.2% FA. Mobile phase B was composed of 70% ACN with 0.2% FA. Samples were separated with a 120 min LC method: peptides were loaded onto the column for 13 min at 0.37 μL/min. Mobile phase B increased from 0 to 6% in 13 min, then to 53% B at 104 min, 100% B at 105 min and held for 4 min at 100% B, decreased to 0% B at 110 min, and a 10 min re-equilibration at 0% B.

Eluting peptide fragments were ionized by electrospray ionization and analyzed on a Thermo Orbitrap Eclipse. Survey scans of precursors were taken from 300 to 1350 m/z at 240L000 resolution. Maximum injection time was 50 ms and automatic gain control (AGC) target was 1E6 ions. Tandem MS was performed using an isolation window of 0.5 Th with a dynamic exclusion time of 10 s. Selected precursors were fragmented using a normalized collision energy level of 25%. MS2 AGC target was set at 2E4 ions with a maximum injection time of 14 ms. Scan range was 150-1350 m/z. Scans were taken at the Turbo speed setting and only peptides with a charge state of +2 or greater were selected for fragmentation.

#### Lipidomics

Extracted lipids were separated on an Acquity CSH C18 column (100 mm x 2.1 mm x 1.7 µm particle size; Waters) at 50°C using the following gradient: 2% mobile phase B from 0-2 min, increased to 30% B over next 1 min, increased to 50% B over next 1 min, increased to 85% over next 14 min, increased to 99% B over next 1 min, then held at 99% B for next 7 min (400 µL/min flow rate). Column re-equilibration of 2% B for 1.75 min occurred between samples. For each analysis 10 µL/sample was injected by autosampler. Mobile phase A consisted of 10 mM ammonium acetate in 70:30 (v/v) acetonitrile:milliQ H2O with 250 µL/L acetic acid. Mobile phase B consisted of 10 mM ammonium acetate in 90:10 (v/v) isopropanol:ACN with 250 µL/L acetic acid.

The LC system (Vanquish Binary Pump, Thermo Scientific) was coupled to a Q Exactive Orbitrap mass spectrometer through a heated electrospray ionization (HESI II) source (Thermo Scientific). Source and capillary temperatures were 300°C, sheath gas flow rate was 25 units, aux gas flow rate was 15 units, sweep gas flow rate was 5 units, spray voltage was |3.5 kV| for both positive and negative modes, and S-lens RF was 90.0 units. The MS was operated in a polarity switching mode; with alternating positive and negative full scan MS and MS2 (Top 2). Full scan MS were acquired at 17,500 resolution with 1 x 10^6^ AGC target, max ion accumulation time of 100 ms, and a scan range of 200-1600 m/z. MS2 scans were acquired at 17,500 resolution with 1 x 10^5^ AGC target, max ion accumulation time of 50 ms, 1.0 m/z isolation window, stepped normalized collision energy (NCE) at 20, 30, 40, and a 10.0 s dynamic exclusion.

The LC system (Vanquish Binary Pump, Thermo Scientific) was coupled to a Q Exactive HF Orbitrap mass spectrometer through a heated electrospray ionization (HESI II) source (Thermo Scientific). Source and capillary temperatures were 350°C, sheath gas flow rate was 45 units, aux gas flow rate was 15 units, sweep gas flow rate was 1 unit, spray voltage was 3.0 kV for both positive and negative modes, and S-lens RF was 50.0 units. The MS was operated in a polarity switching mode; with alternating positive and negative full scan MS and MS2 (Top 10). Full scan MS were acquired at 60K resolution with 1 x 10^6^ AGC target, max ion accumulation time of 100 ms, and a scan range of 70-900 m/z. MS2 scans were acquired at 45K resolution with 1 x 10^5^ AGC target, max ion accumulation time of 100 ms, 1.0 m/z isolation window, stepped NCE at 20, 30, 40, and a 30.0 s dynamic exclusion.

### Data analysis for proteomics and lipidomics

#### Proteomics

LC-MS files for proteomics were searched in Maxquant (version 2.0.3.1). Original outputs from Maxquant were inspected and potential contaminant proteins, protein groups that contain proteins identified with decoy peptide sequence, and those identified only with modification site were removed. LFQ intensities were used as the quantification metric.

#### Lipidomics

LC-MS files for lipidomics were processed using Compound Discoverer 3.1 (Thermo Scientific) and LipiDex (Hutchins et al., 2018). All peaks with a 1.4-23 min retention time and 100 Da to 5000 Da MS1 precursor mass were aggregated into compound groups using a 10 ppm mass tolerance and 0.4 retention time tolerance. Peaks were excluded if peak intensity was less than 2 x 106, peak width was greater than 0.75 min, signal-to-noise ratio was less than 1.5, or intensity was < 3-fold greater than blank. MS2 spectra were searched against an in-silico generated spectral library containing 35,000 unique molecular compositions of 48 distinct lipid classes (Hutchins et al., 2019). Spectra matches with a dot product score > 500 and reverse dot product score > 700 were retained for further analysis. Lipid MS/MS spectra that contained < 75% interference from co-eluting isobaric lipids, eluted within a 3.5 median absolute retention time deviation (M.A.D. RT) of each other, and were found within at least 4 processed files were used for identification at the individual fatty acid substituent levels of structural resolution. If individual fatty acid substituents were unresolved, then identifications were made with the sum of the fatty acid substituents. Peak intensities were normalized with the peptide amount to correct for different amounts of starting materials across the RIAIL panel.

### Survival analysis and lifespan traits extraction

The *survfit* function of the survival (version 3.5-0) R package was used to analyze survival data. The following formula was used “*survival::Surv(Age_of_death, status) ∼ Strain*” with default parameters. Parental strains (N2 and CB4856) lifespan from each batch was compared to check for possible batch effects. No batch correction was performed. The *quantile* function was used to obtain the average lifespan as well as the 25%, 50% and 75% mortality.

### Life-history trait batch correction

The *lmer* function of the lmerTest (version 3.1-3) R package was used to adjust for batch effects in data collection. The following formula was used “*value ∼ (1|batch/channel)*” with default parameters.

### Statistical analyses

In the analysis of continuous variables across groups, we computed p-values using two-sided Student’s t-tests to ascertain statistical significance (Figures 5B, 5C, and S6C). To explore relationships among variables we used Pearson correlations (Figures 2F, 2G, 3F, and S8A). Resulting p-values (where applicable) were corrected for multiple testing using the Benjamini–Hochberg false discovery rate.

### Dirichlet regression analysis

The DirichletReg (version 0.7-1) R package was used to analyze the shape proportion data and generate visualizations (Figures 2H and S2). Univariate analysis among variables was performed with default parameters following package documentation.

### Variant calling and genetic map construction

Using the RNA-seq data, we performed variant calling employing the Genome Analysis Toolkit (GATK) (version 4.2.4.0) (Brouard et al., 2019) following their best practices workflow (Figure S7) (Van der Auwera et al., 2013) to genotype the 85 RIAILs strains. In brief, RNA-seq reads were mapped to the reference genome using STAR and prepared for variant calling (Mark Duplicates, SplitNCigarReads, Base Quality Recalibration). Then short variants (SNPs and Indels) were called using GATK’s HaplotypeCaller. Next, we exploited the design of the study (parental replicates and strain under control and treated conditions) to obtain a high-confidence set of germline variants. Comparison of identified variants with publicly available variant information for *C. elegans* (https://www.elegansvariation.org/) and previous genetic work on the RIAILs allowed us to perform quality control checks on the obtained variants (Figures S7 and S8). We then used the onemap (version 2.8.2) (Margarido et al., 2007) software to construct a genetic map for subsequent QTL mapping. For comparison to known variants (Figure S7B), variants for the CB4856 strain were retrieved from the “Caenorhabditis elegans Natural Diversity Resource” (https://elegansvariation.org/). Soft filtered variant file was retrieved from http://storage.googleapis.com/elegansvariation.org/releases/20210121/variation/WI.20210121.hard-filter.vcf.gz. Hard filtered variant file was retrieved from http://storage.googleapis.com/elegansvariation.org/releases/20210121/variation/WI.20210121.soft-filter.vcf.gz. These variants were then filtered for the CB4856 strain keeping only 1/1 variants with high impact consequence.

### Association mapping and gene-set enrichment analysis

We used the *lmekin* function of the coxme R package as it allows one to model the correlation structure of the random effects. Analysis of deviance for *lmekin* from https://aeolister.wordpress.com/2016/07/07/likelihood-ratio-test-for-lmekin/ was used to calculate Likelihood ratio between the null model “*value ∼ 1 + (1|kinshipCov)*” and the model of interest “*value ∼ 1 + predictorValue + (1|kinshipCov)*”. An INT transformation was to transform the data before mapping. The Benjamini-Hochberg procedure was selected for multiple-testing correction. As the traits mapped are not independent, such a correction would be over-correcting. GSEA analysis was performed using Clusterprofiler (version 4.2.2) and org.Ce.eg.db (version 3.14.0). The used gene list was ranked by the signed LOD-value obtained from the association mapping analysis.

### Lipid class over-representation analysis

All measured lipid species were used to define lipid class sets. These were used along with the *enricher* function from the clusterProfiler (version 4.2.2) R package to conduct lipid class enrichment analysis (Figure S5), which is designed to accept customized annotations through the TERM2GENE parameter. The Benjamini-Hochberg procedure was selected for multiple-testing correction. Enrichment was tested for each lifespan trait (average lifespan, 25%, 50% and 75% mortality) for two groups of lipids: positively associated (non-adjusted p-value < 0.05 & association coefficient > 0) and negatively associated (non-adjusted p-value < 0.05 & association coefficient < 0).

### Quantitative trait locus (QTL) mapping

The qtl2 (version 0.34) R package (Broman et al., 2019) was used to perform QTL mapping of all phenotypic and molecular traits. An INT transformation was used to transform the data before mapping. Gene codes were encoded as N2 = 1, CB4856 = 2 and heterozygotic (N2/CB4856) = 3. Crosstype was specified as “risib”. Pseudomarkers were inserted into the genetic map with a step of 1 and default values for other parameters. Conditional genotype probabilities, kinship and genome scans were performed using qtl2 package functions with default parameters. Significance thresholds for each trait were obtained through permutation testing using the *scan1perm* qtl2 function with 1000 permutations. Finally, the *find_peaks* function was used to identify significant QTLs with a threshold of 0.05 and a drop of 0.5.

### Lifespan locus differential analysis and gene-set enrichment analysis

Concerning Figures 5B and 5C, RIAIL strains were split into two groups based on their genotype at position 13,121,591 on Chr. II and 12,125,475 on Chr. V. Using the package limma (version 3.50.1) we performed differential expression analysis between the two groups of RIAILs. GSEA analysis was performed using Clusterprofiler (version 4.2.2) and org.Ce.eg.db (version 3.14.0). The Benjamini-Hochberg procedure was selected for multiple-testing correction. The used gene list was ranked by the signed p-value obtained from the differential expression analysis.

### UK Biobank GFM1 and RICTOR SNP-disease time-to-event analysis

The time-to-event analysis was performed in the UK Biobank, a population cohort of ∼500,000 participants from the United Kingdom (Sudlow et al., 2015) (project 48020). The sample analyzed was restricted to participants of European ancestry (as determined in Pan-UKBB, https://pan.ukbb.broadinstitute.org) who were unrelated, as determined by their inclusion in the original calculation of the genetic principal components (field 22020). Time-to-event was measured from birth to the first occurrence of the event. We selected 19 diseases to include as events in addition to death, listed in the table below. Variants were selected from whole-exome sequencing, where at least 5 minor alleles were detected. These selection criteria resulted in 339’967 individuals and 821 and 577 SNPs for *GFM1* and *RICTOR*, respectively.

The time-to-event analysis was done with Cox proportional hazards in R using the Coxph function from the survival packages (Borgan, 2001; Therneau, 2023). The top 40 genetic principal components, sex, and the batch (specifically the initial 50k released, field 32050) were included as covariates. In some cases, the maximum likelihood of the method failed to converge, generally due to no events being recorded in individuals with alternate alleles, in which case the results were marked as unreliable for that SNP-event combination. These were retained only for the purpose of multiple-testing correction, which was performed using the Benjamini-Hochberg method (Benjamini and Hochberg, 1995).

| **Event** | **ICD10 code(s)** | **Number of events** |
| --- | --- | --- |
| Myocardial infarction | I21, I22, I24 | 18,016 |
| Cardiomyopathy | I42 | 2,584 |
| Heart failure | I50 | 14,097 |
| Chronic ischaemic heart disease | I25 | 37,124 |
| Aortic aneurysm and dissection | I71 | 3,733 |
| Cerebrovascular disease | G45, I60-I64, I67, I68 | 23,710 |
| Vascular dementia | F01 | 1,493 |
| Parkinson disease | G20, G23 | 3,228 |
| Secondary parkinsonism | G21 | 226 |
| Alzheimer's disease/dementia | F00, F02, F03, G30 | 5,582 |
| Multiple sclerosis and other demyelinating diseases of the central nervous system | G35, G37 | 2,036 |
| Insulin-dependent diabetes mellitus | E10 | 3420 |
| Non-insulin-dependent diabetes mellitus | E11, E13, E14 | 31,663 |
| Acute pancreatitis | K85 | 3,391 |
| Acute renal failure | N17 | 17,241 |
| Chronic renal failure | N18 | 20,369 |
| Alcoholic liver disease | K70 | 1,627 |
| Fibrosis and cirrhosis of the liver | K74 | 2,044 |
| Other inflammatory liver diseases | K75 | 2,242 |
| Death | - | 25,915 |

### Figures and visualizations

Data visualization was performed using ggplot2 (version 3.4.2). The resulting p-values (where applicable) were corrected for multiple testing using the Benjamini–Hochberg false discovery rate. Clusterprofiler (version 4.2.2) was used to generate graph representations of enrichment results (Figures 3D and S4D). The R package enrichplot (version 1.14.2) was used to generate running GSEA plots (Figure 3E). UpSetR (version 1.4.0) was used to generate upset plots (Figures 3C, S3, and S4C).

### Data availability

The RNA-Seq data generated in this study have been deposited in the GEO database (GSE252593). The remaining data generated in this study are provided in the Source Data files. Scripts for analysis and figure generation have been deposited in a GitHub (https://github.com/auwerxlab/Project_RIAILs) repository along with additional data used in this work.

### Fluorescent image for assessing the UPR^mt^ activation

RNAi bacteria were cultured overnight in lysogeny broth (LB) medium containing 100 mg/mL ampicillin at 37°C. Then the bacteria were five times concentrated and seeded onto RNAi plates. Random L4/young adult worms were picked onto the RNAi bacteria-seeded plates and cultured at 20°C until their progenies reached the young adult stage. 6 - 10 worms were then randomly picked in a drop of 20 mM tetramisole (Cat. T1512, Sigma) and then aligned on an empty NGM plate. Fluorescent images, with the same exposure time for each condition, were captured using a Nikon SMZ1000 microscope.

### Real-time quantitative PCR (RT-qPCR)

For qRT-PCR, worms were cultured and total RNA was extracted as described for the RNA-seq sample preparation. cDNA synthesis was performed using the Qiagen Reverse Transcription Kit (205314) from the extracted RNA samples. The qPCR was then conducted with the Roche Light Cycler 480 SYBR Green I Master kit (Cat. 04887352001). The specific primers utilized are detailed in the key resources table, with *pmp-3* primers serving as housekeeping controls.

### OCR measurements by Seahorse

Oxygen consumption rate (OCR) was assessed using the Seahorse XF96 (Seahorse Bioscience), following the protocol outlined in (Koopman et al., 2016). Briefly, a synchronized culture of ∼100 worms was harvested on day 1 of adulthood with sterile M9 buffer. After three washes in the M9 buffer, the worms were transferred to a 96-well Seahorse plate, where their OCR was measured six times to determine mitochondrial activity for each condition at basal level and another six times measurement after adding 10 µM FCCP as the final concentration.

## QUANTIFICATION AND STATISTICAL ANALYSIS

No statistical methods were applied to pre-determine worm sample size. Comparison between more than two groups was assessed by using a One-way ANOVA test. Prism 8 (GraphPad Software) was used for statistical analysis of all lifespan, qRT-PCR, OCR, and paralysis experiments. Variability in panels is given as the s.e.m. All p<0.05 were considered to be significant. (\*\*\*\**p*<0.0001; \*\*\**p*<0.001; \*\**p*<0.01; \**p*<0.05; n.s., not significant. For lifespan, and OCR measurement in worms, sample size was determined based on the known variability of the experiments. All experiments were done non-blinded.

