## Supplemental Figures for "High-content phenotypic analysis of *a C. elegans* recombinant inbred population identifies genetic and molecular regulators of lifespan"

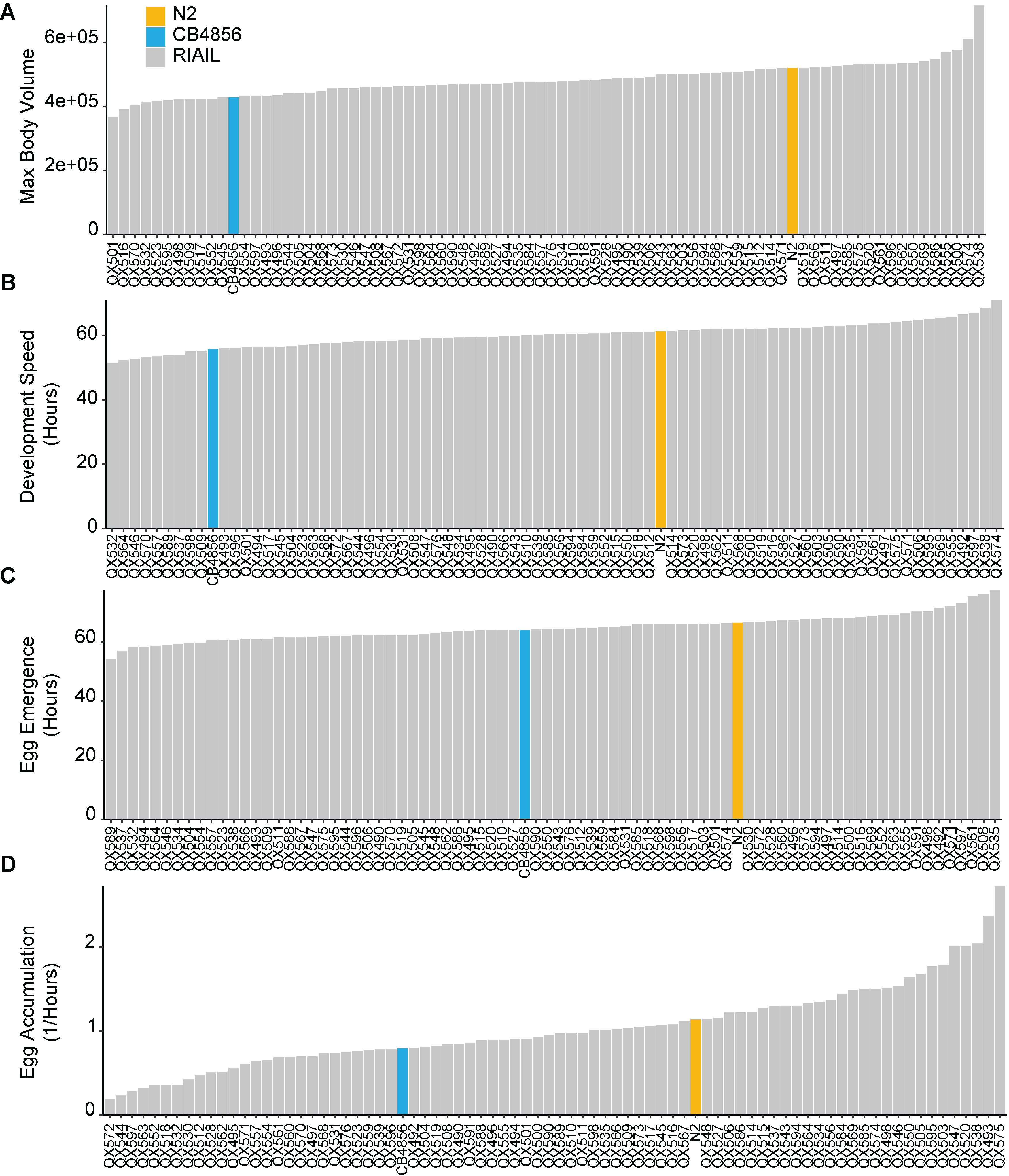

**Supplemental Figure S1. The RIAIL life-history traits**. Barplots showing life-history traits of 85 RIAIL strains and two wild-type parental strains. Grey bars: RIAILs; orange bars: N2 (Bristol); blue bars: CB4856 (Hawaii, HW) strains for maximal body size (**A**), developmental time (**B**), time of the first egg (**C**), and speed of egg accumulation (**D**). The Y-axis value represents scaled residuals following the removal of batch and channel effects. **Related to Figure 2.**

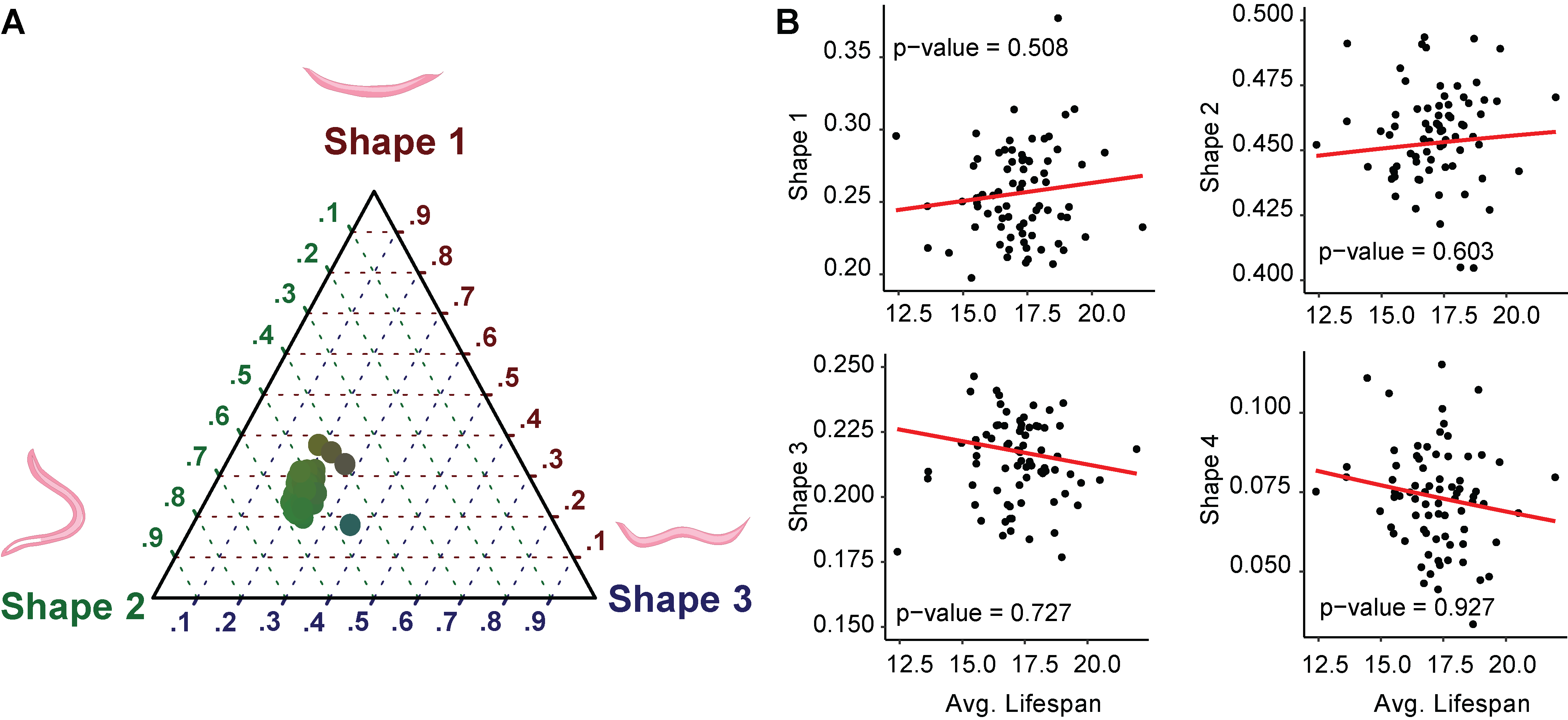

**Supplemental Figure S2. No correlations between four moving shapes and the average lifespan of RIAILs.** (**A**) Ternary plot of the proportion spent between 4 different shapes during the observation period (egg to gravid adults). Shape 1: molting or dead; Shape 2: active; Shape 3: swimming. (**B**) Scatter plot of the average lifespan of each RIAIL strain against the proportion of time spent in different shapes. The red line represents the fit estimate for each shape calculated using the *Dirichletreg* package. **Related to Figure 2.**

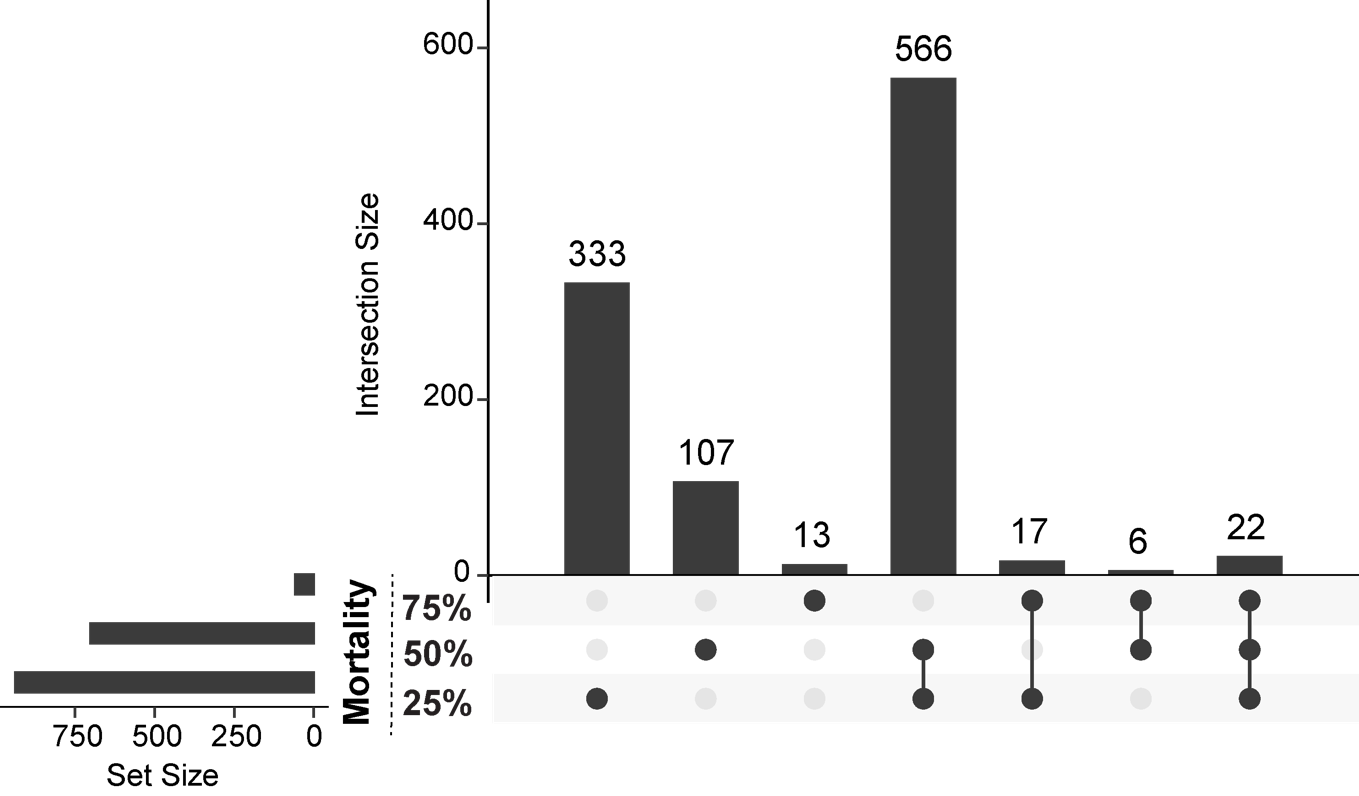

**Supplemental Figure S3.** **Upset plot of significantly enriched (adjusted p-value < 0.05) biological process genesets of mRNA-lifespan associations.** The number of genesets unique or shared between different categories is indicated on top of the bars. Lifespan traits are ranked based on the number of associations. **Related to Figure 3 and Tables S1 and S2.**

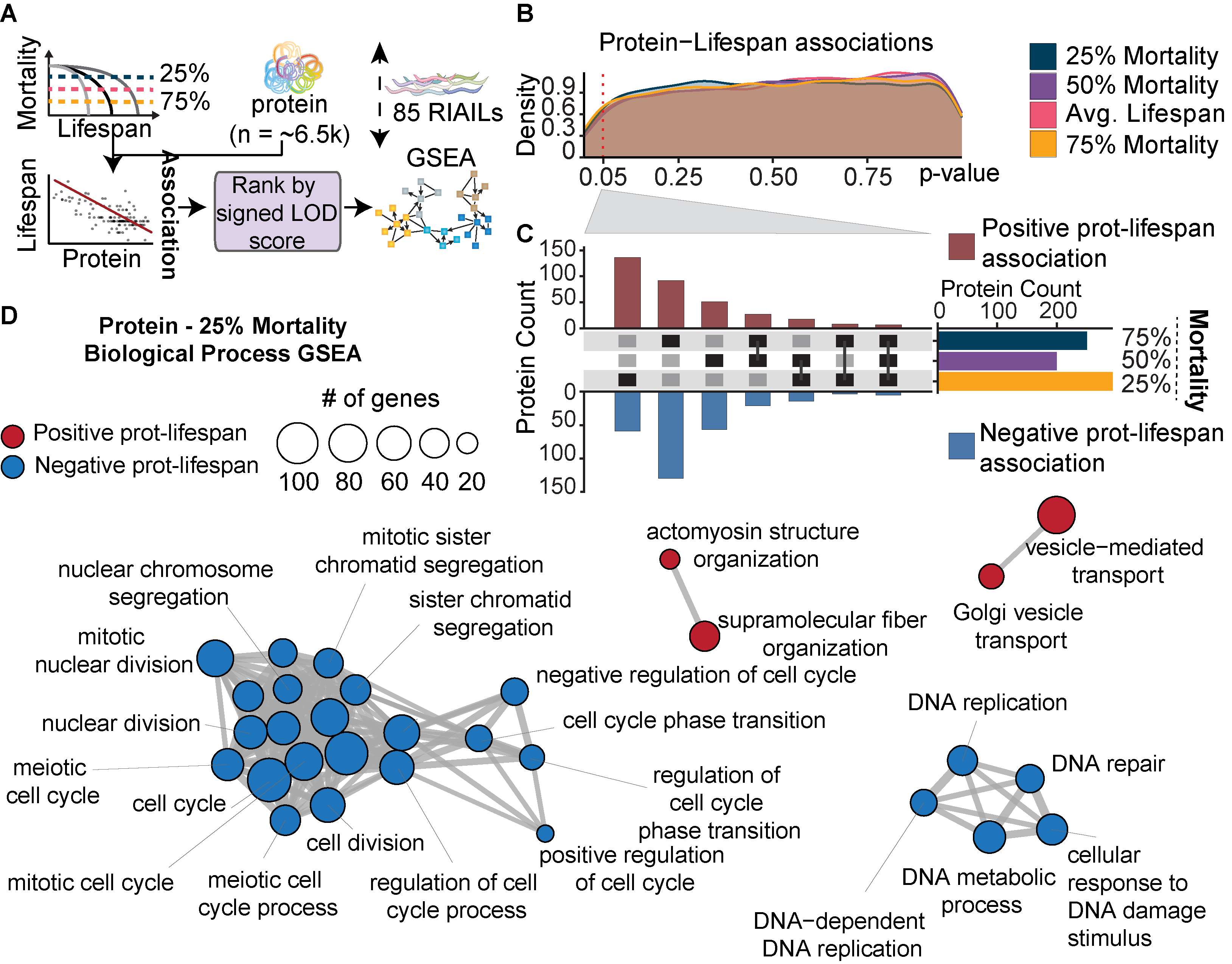

**Supplemental Figure S4.** **Quantitative assessment of proteome-lifespan associations.** (**A**) Diagram of protein-lifespan association mapping and geneset enrichment analysis (GSEA). (**B**) Line histogram of non-adjusted p-values of protein-lifespan associations for the different lifespan traits. The Y-axis represents the density of p-values. The red dashed vertical line represents a p-value of 0.05. Color represents the different lifespan traits. (**C**) The upset plot of positively- (red) and negatively- (blue) associated proteins with 25%, 50%, and 75% mortality in the RIAILs with a non-adjusted p-value < 0.05. (**D**) Graph representing top 30 biological process genesets enriched. Geneset enrichment analysis of protein-25% mortality associations. Genes of proteins were ranked by the signed logarithm of the odds (LOD) score of protein-25% mortality association. Color represents positive (red) or negative (blue) normalized enrichment score (NES). All genesets in the figure had a significant q-value. Genesets in bold: overlapping genesets in both mRNA and protein levels. **Related to Table S3.**

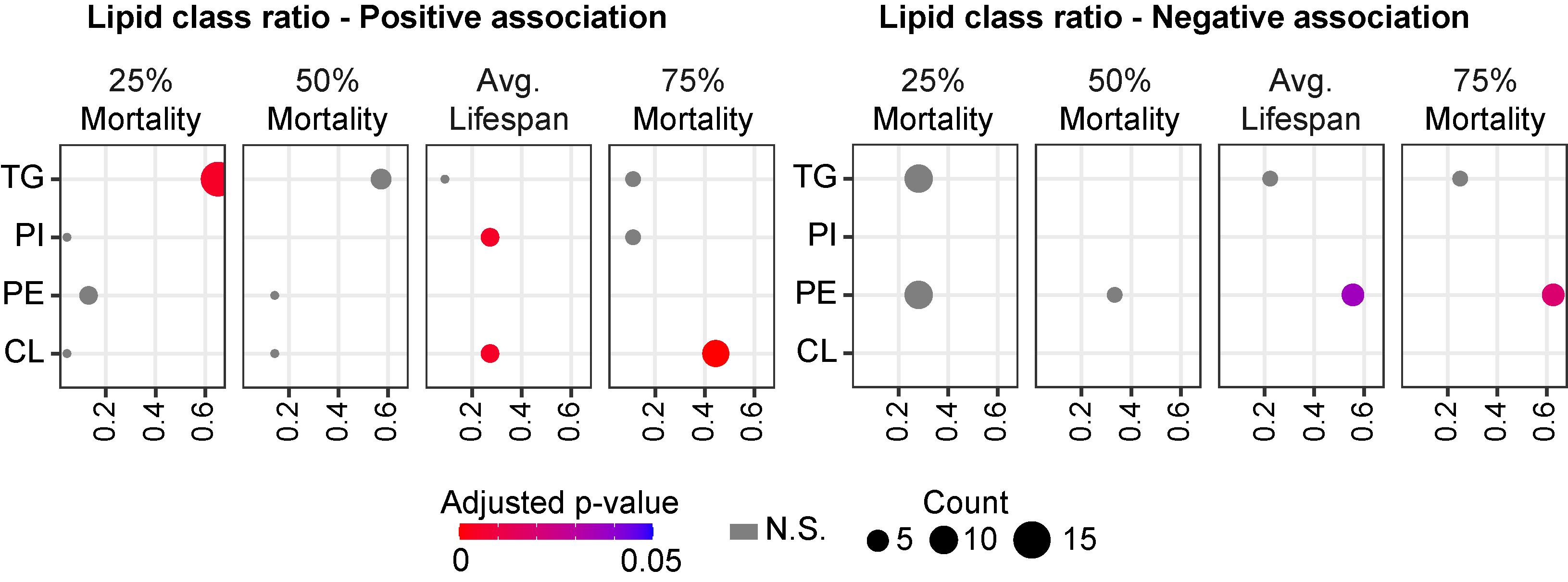

**Supplemental Figure S5. Over-representation of lipid class in significant lipid-lifespan association results for each lifespan trait.** TG: triglycerides. PI: phosphatidylinositol. PE: phosphatidylethanolamine. CL: cardiolipin. Color represents adjusted p-values. Dot size represents the number of lipids (lipid count). N.S.: not significant. **Related to Figure 4 and Table S4.**

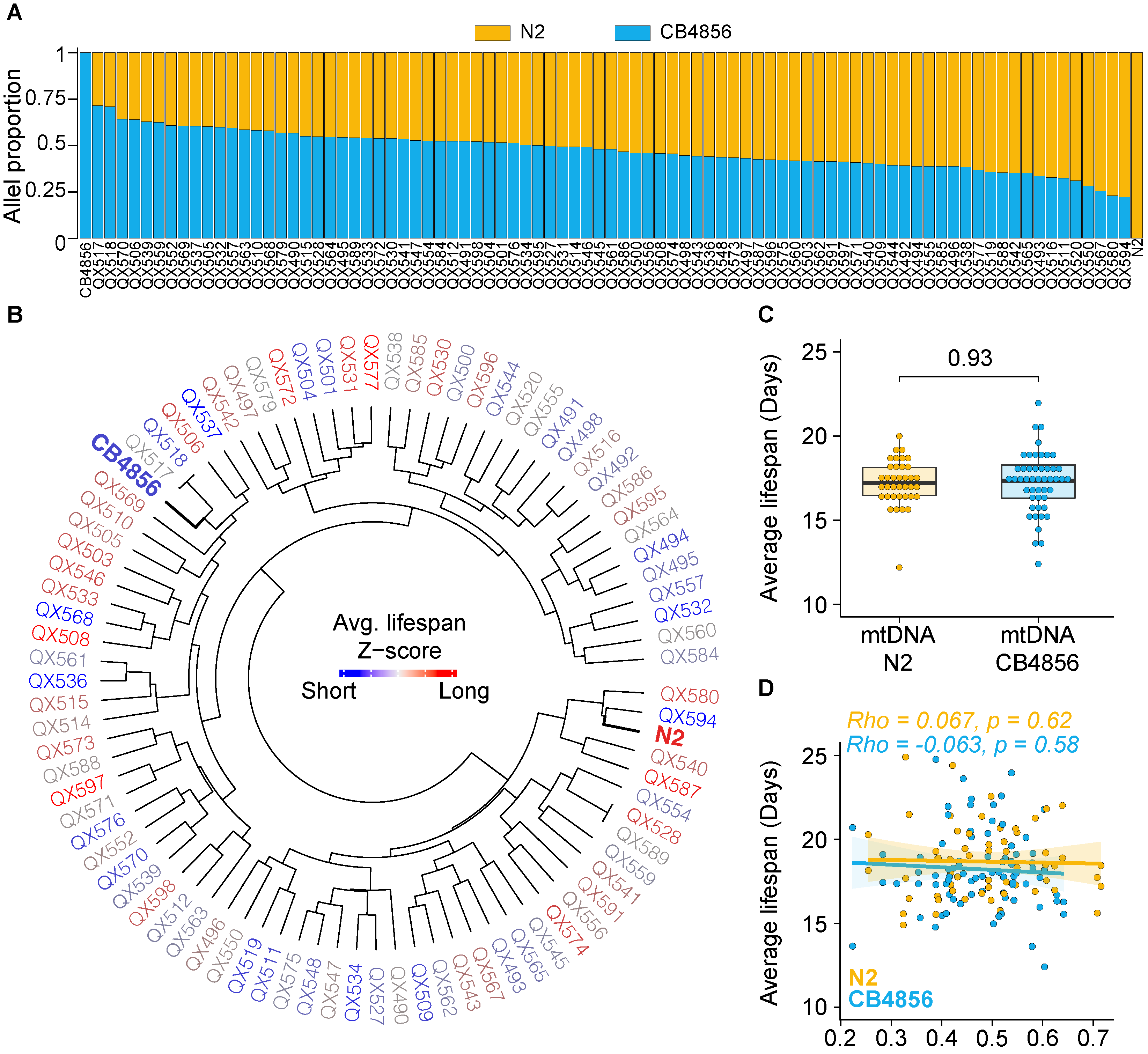

**Supplemental Figure S6. RIAIL Lifespan is not simply explained by genomic composition and mitochondrial haplotype.** (**A**) Stacked barplot of Hawaii allele content (HAC) of CB4856 and N2 of 85 RIAIL strains and two wild-type parental strains. (**B**) Phylogenetic tree of the 85 RIAIL strains and the 2 parental strains (N2 and CB4856, in bold) using the overall kinship based on all identified variants. Color represents the z-score of the average lifespan for each strain. (**C**) Boxplot of the average lifespan of RIAILs strains with CB4856 or N2 mitochondrial DNA at control conditions. The p-value represents the comparison of the two groups calculated using a two-tailed Student’s t-test. (**D**) The scatter plot of HAC and average lifespan of each RIAIL strain. Color represents strains with N2 (orange) or CB4856 (blue) mitochondrial DNA. Spearman correlation coefficients with p-values.

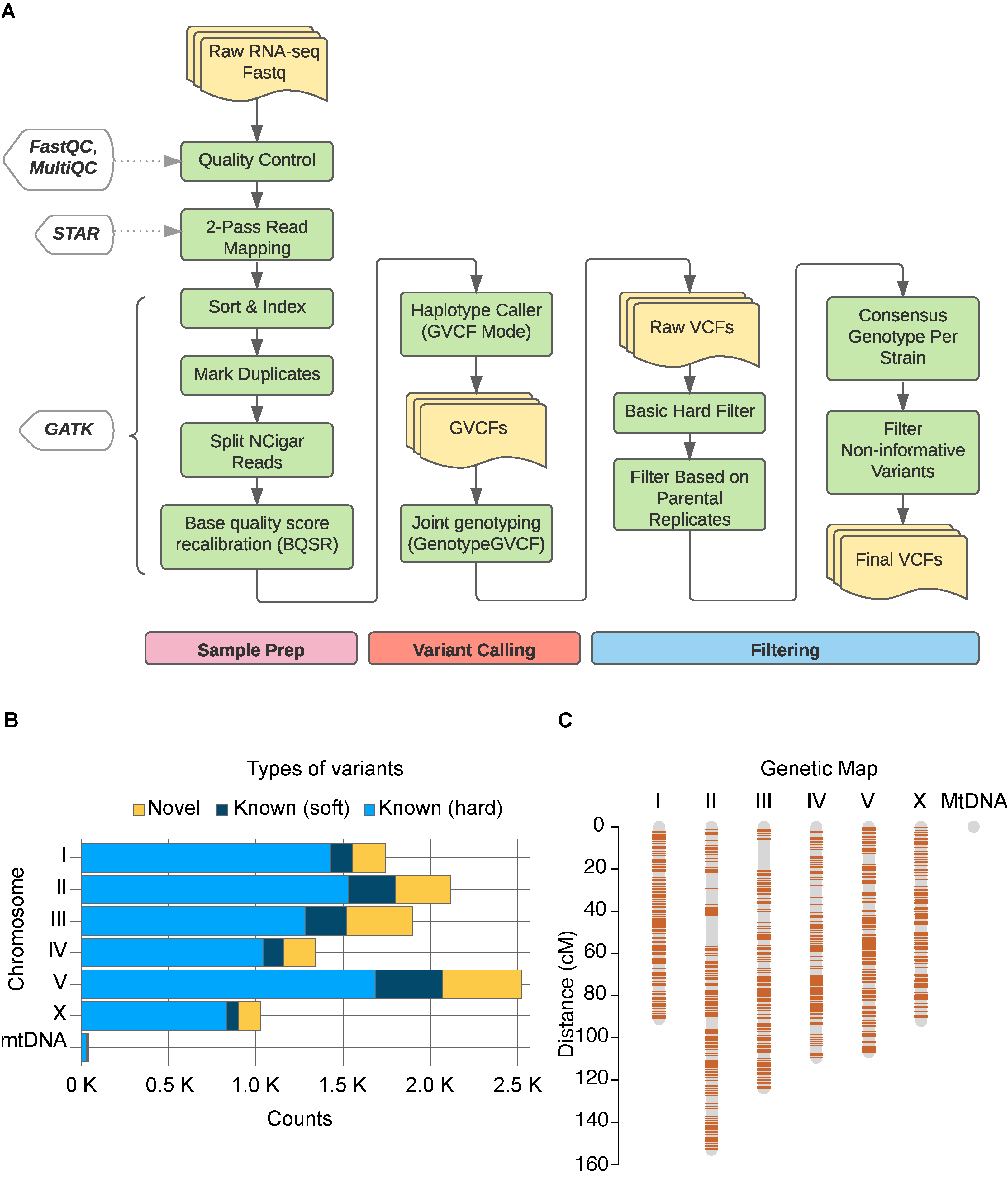

**Supplemental Figure S7.** **Variant calling and genetic map building of the RIAILs from RNA-seq data**. (**A**) The pipeline for sample preparation, variant calling, and filtering. Detailed procedures are provided in the methods. (**B**) Comparison of identified variants with publicly available variant information for *C. elegans* (<https://www.elegansvariation.org/>). Light blue: known variants based on a hard filter. Dark blue: known variants based on a soft filter. Yellow: novel variants. (**C**) Generated RIAIL genetic map using identified variants. **Related to Figure 5.**

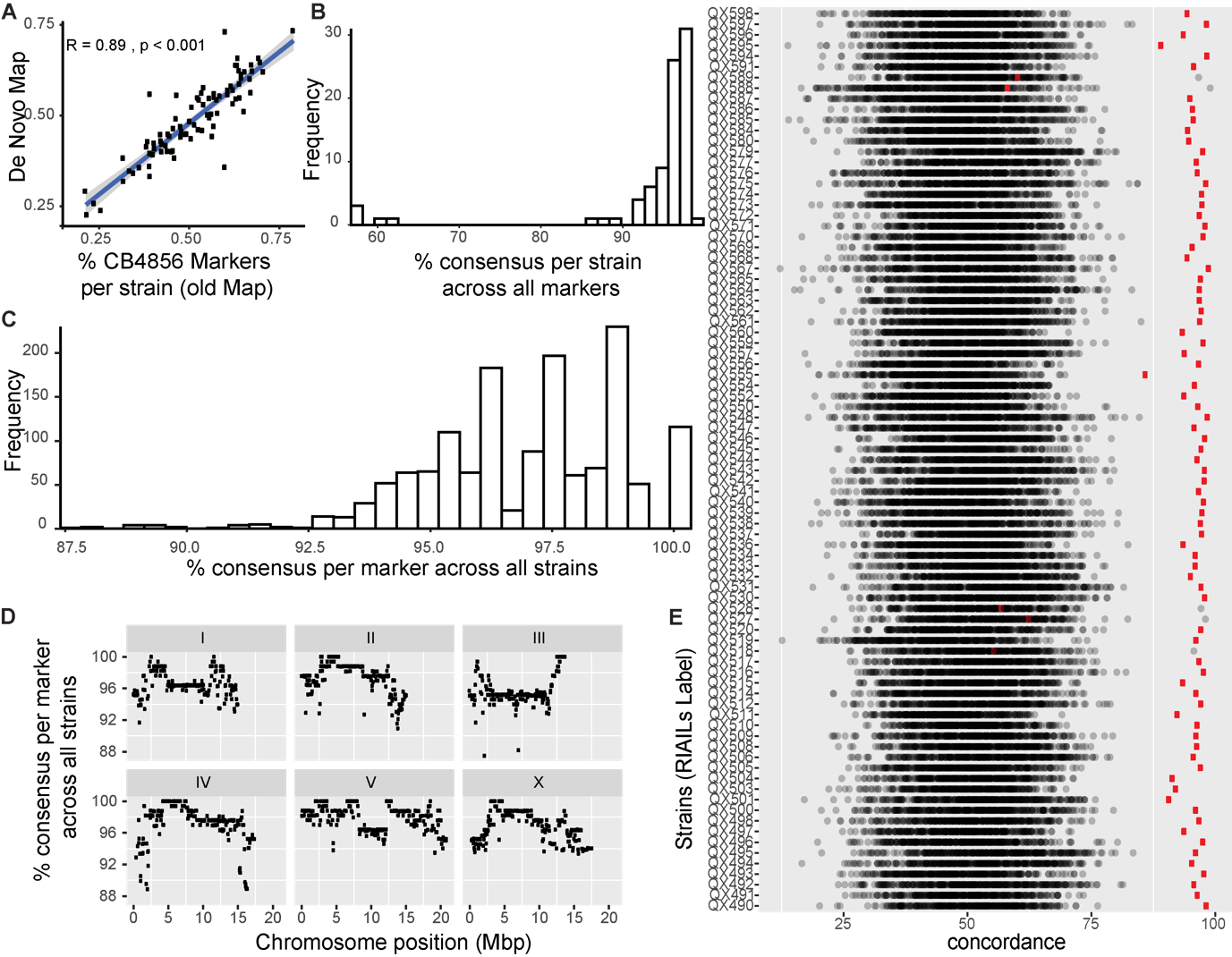

**Supplemental Figure S8.** **Comparison of *de novo* RIAILs map to legacy map (Andersen et al., 2015)**. (**A**) Scatter plot of percentage of Hawaiian strain (CB4856) markers per strain in the legacy map (x-axis) and *de novo* map (y-axis). Pearson correlation coefficient with p-value. (**B**) Histogram of the percent consensus across all markers per strain. The Y-axis represents the number of strains with a given % consensus (x-axis). (**C**) Histogram of the percent consensus across all strains per marker. The Y-axis represents the number of markers with a given % consensus (x-axis). (**D**) Scatter plot of the percent consensus across all strains per marker (y-axis) vs the chromosomal position (x-axis) per chromosome in megabase pairs (Mbp). (**E**) Plot of the concordance between each RIAIL strain and all other strains to check for possible miss-labeled strains. Concordance is calculated as the percentage of identical markers between the *de novo* map for that strain and the legacy map for each of the other strains. The red mark represents the concordance value for the strain itself. **Related to Figure 5.**

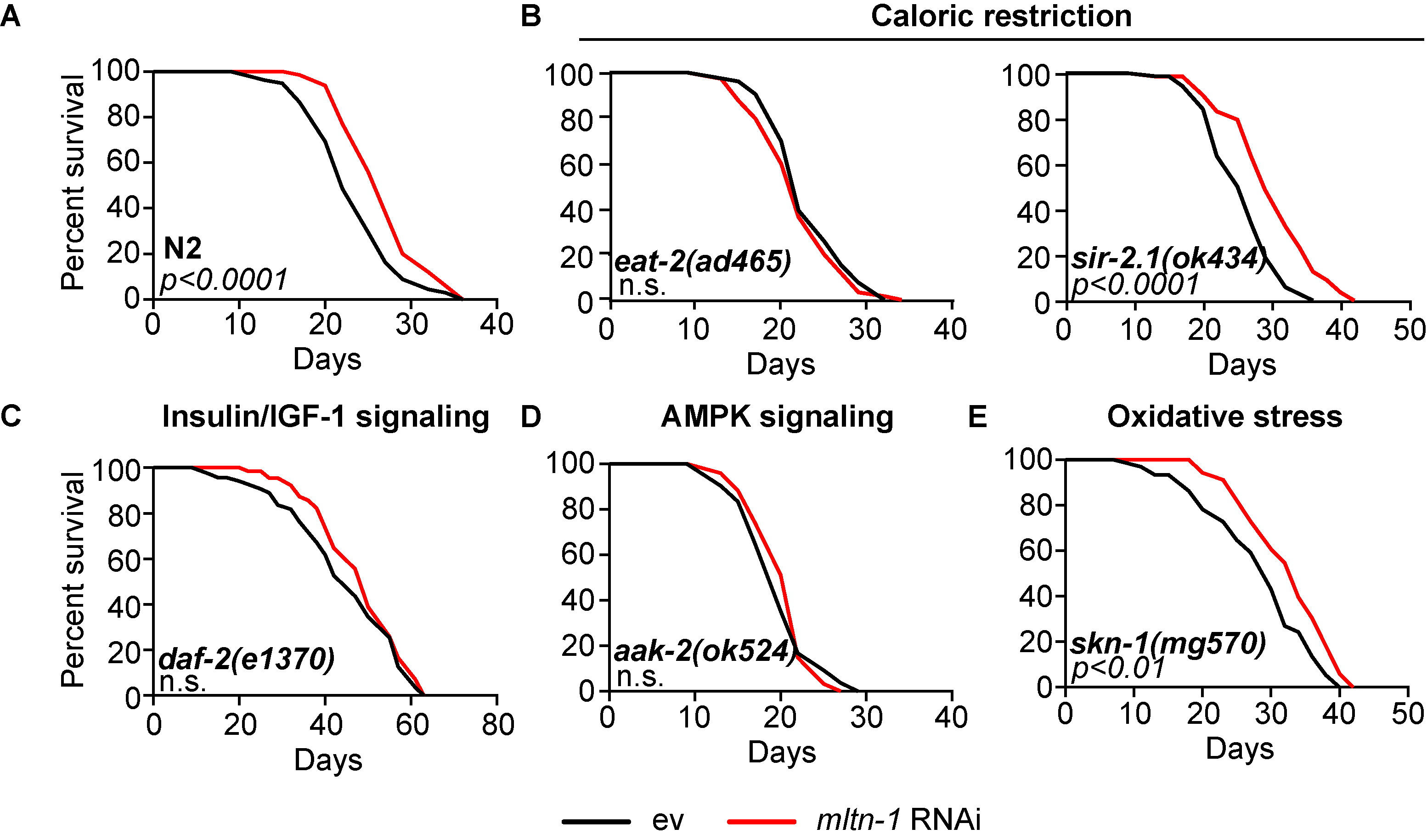

**Supplemental Figure S9. Validation of *mltn-1* RNAi in known longevity pathways.** (**A**) Confirmation of lifespan extension observed in N2 worms fed with *mltn-1* RNAi. (**B**) Lifespan measurement of *mltn-1* RNAi in caloric restriction models, such as *eat-2(ad465)* and *sir-2.1(ok434)* worms. (**C-E**) Lifespan measurements of *daf-2(e1370)*(insulin/IGF-1 signaling), *aak-2(ok524)*(AMPK signaling), and *skn-1(mg570)*(oxidative stress) worms fed with either control or *mltn-1* RNAi. **Related to Figure 6.**
